# Mining and e-waste recycling influence the spatial distribution of technology-critical elements, but not rare earth elements, in boreal lakes

**DOI:** 10.1101/2025.09.02.673822

**Authors:** Marta Gabriele, Maikel Rosabal, Miguel Montoro Girona, Patrice Blaney, Guillaume Grosbois

**Author notes:** Corresponding author: Marta Gabriele.

## Abstract

Mining and more recent e-waste recycling have contributed trace elements (TEs) to the environment. However, the occurrence of emerging technology-critical elements (TCEs), including rare earth elements (REEs), remains poorly reported. Our study aims to i) investigate the spatial distribution of TEs, including TCEs, across different environmental matrices; ii) compare measured concentrations in water and sediment against environmental quality guidelines; and iii) assess potential risks to human health from fish consumption. In this study, we sampled water, sediment, and fish tissues (muscle and liver) across six boreal lakes near the historically mining region of Rouyn-Noranda, home to North America’s largest copper smelting and recycling facility (Horne Smelter). Concentrations of TEs (e.g., Cu, Se) were higher in lakes closest to the smelter. Similarly, some TCEs (i.e., Ti, Co, Tl) followed this same spatial distribution pattern, suggesting that their release may be linked to historical and current mining activities. Conversely, REEs displayed distinct spatial patterns, likely influenced by geological sources rather than pollution. Several TEs (e.g., Zn, Cd, Pb) exceeded Canadian water and sediment quality guidelines in lakes closer to the mining area. Muscle tissue from walleye or yellow perch showed Zn, Cd, or Pb concentrations above safety limits in at least one lake. This study highlights the importance of including emerging TCEs (e.g., Sr, Tl, Co) in biomonitoring programs. Our findings provide critical insights into the environmental distribution of TEs across multiple matrices of boreal lake ecosystems, contributing to global efforts in risk assessment and sustainable freshwater management in the context of growing electronic waste recycling.

**Highlights:**

▪ First report of various TCEs and REEs in 3 matrices of boreal lakes
▪ The spatial distribution of several newly reported TCEs is similar to that of historical TEs
▪ REE concentrations are associated with geology rather than mining sources
▪ Some TE concentrations in water and sediment are above guidelines in closest lakes
▪ All lakes exceeded safety limits for at least one element (Zn, Cd, or Pb) in the muscle tissue of either walleye or yellow perch

## 1. Introduction

Freshwater ecosystems, which cover about 0.8% of the Earth’s surface provide essential ecosystem services, such as fish production, drinking water supply, recreation, and climate regulation (Lynch et al., 2023). However, over the past two centuries, human demographic growth has substantially increased anthropogenic pressures on these ecosystems by driving land conservation and pollution (Bowler et al., 2020). Among these pressures, mining stands as one of the top five threats to biodiversity loss (Dudgeon et al., 2006). Besides altering natural habitats, it generates large amounts of waste that enter rivers and lakes, contributing to ecosystem degradation at both local and global scales (Salomons et al., 1987).

Among the pollutants released, trace elements (TEs)—metals (e.g. Cu, Cd, Zn) and metalloids (e.g. As, Sb)—pose major threats to human health (Thompson and Darwish, 2019) and to aquatic organisms (Le Saux et al., 2020). Their presence and distribution are influenced by both natural (e.g., erosion of the continental crust) and anthropogenic (e.g., mining) processes, which can lead to excessive accumulation in different environmental matrices such as water, sediment, and biota, including fish (Ali et al., 2021; Taylor and McLennan, 1995). Metal accumulated in edible fish tissues presents significant health risks to consumers (Sow et al., 2013) once their levels exceed safety thresholds. Additionally, high concentrations of TEs in aquatic ecosystems can also adversely affect resident organisms (Garai et al., 2021; Ghosh et al., 2022) (e.g., fish communities), elevating mortality rates and reducing populations growth and abundance (Blaney et al., in review; Mayer-Pinto et al., 2010). Thus, to assess the environmental impacts of industrial activities and to ensure restoration of degraded areas, we must understand the environmental behavior of TEs across waterbodies and organisms.

Certain TEs such as Ti, Co, Tl, Sr, Sb, and rare earth elements (REEs) (Willner et al., 2021; Trimmel et al., 2023) have become increasingly in demand to produce emerging high technologies, including cell phones, laptops, photovoltaic cells, and semiconductors (Cenci et al., 2024). As their use in modern society rises, these elements—referred to as technology critical elements (TCEs)—become essential components of human daily life. However, the lifecycle of these high technologies also amplifies the risk of aquatic contamination as their production involves direct smelter emissions, fossil fuel combustion, and their final disposal of electronic products by recycling activities (Pal et al., 2010; Belzile and Chen, 2017). Hence, a deeper understanding of the distribution and environmental impact of TCEs is crucial, as current knowledge on their impacts remains limited.

The Abitibi-Témiscamingue region contains over 20,000 lakes that support diverse aquatic fauna and flora, including waterbirds, insects, and mammals (Beaulne et al., 2012; Hasan et al., 2023). However, these ecosystems are affected by anthropogenic activities such as forestry and pollution from the multiple industrial and human activities active in the region (Grosbois et al., 2023; Guimond et al., 2024). The city of Rouyn-Noranda, a key industrial hub in Canada, is home to the Horne Smelter (HS) since 1920, the only operating copper (Cu) smelter in Canada (Leverington and Schindler, 2018). The discovery of Cu and gold (Au) deposits in this region spurred rapid economic expansion and intensive extraction activities. Since 1986, the HS has processed electronic waste from Canada and the United States to give Cu and other metals a second life, making it North America’s first metal recycling facility (Telmer et al., 2006). The region has a documented history of contamination by conventional TEs such as Cu, As, Pb, and Zn in both abiotic and biotic samples, such as water, sediments, and fish (Campbell et al., 2003; Darricau et al., 2021; Giguère et al., 2005; Kraemer et al., 2008). However, the shift towards a recycling facility may potentially introduce less frequently monitored elements, including Ti, Co, Sr, Sb, Tl, and REEs. Understanding the spatial distribution of TEs, including TCEs, in lake sediments and waters is essential, as it determines not only the extent of pollution but also the bioavailability and potential uptake by aquatic organisms (Adams et al., 2020). Moreover, mapping how these elements disperse from point sources into surrounding ecosystems provides critical insights into their environmental behavior and allows for distinguishing between natural and anthropogenic sources (Zheng et al., 2008). Recent studies have only begun to reveal spatial variations in the distribution of REEs such as Ce and La in lake water and lichens (Ricard-Henderson, 2023; Dupont et al., 2025), underscoring the need for comprehensive spatial assessments of these emerging contaminants.

This study aims to i) quantify TEs, including emerging TCEs and REEs, in water, sediment, and fish tissues (liver and muscle) across a spatial gradient of lakes, from proximal to distal sites, while considering prevailing wind patterns; ii) compare measured concentrations in water and sediment to environmental quality guidelines; iii) assess potential human health risks associated with fish consumption based on contaminant levels in muscle tissues. We hypothesized that i) the spatial distribution of both historically TEs and newly reported TCEs and REEs would be primarily driven by historical mining activities and atmospheric emissions from the HS given their persistent deposition in surrounding ecosystems through air and water pathways over decades; ii) lakes located closer to the HS would exceed environmental quality guidelines more frequently than farther ones, reflecting higher contamination levels due to their greater proximity to the pollution source; iii) fish from distant lakes with higher contamination are expected to exhibit higher concentrations of TEs (Cu, As, Zn, Pb) due to bioaccumulation from water and sediments, potentially exceeding thresholds for safe human consumption and indicating heighten risk to populations dependent on these resources.

## 2. Materials and methods

### 2.1. Study area and lake selection

The study sites are located around the city of Rouyn-Noranda, in the Abitibi-Témiscamingue region of Québec, Canada (Fig. 1). Lakes in this region host numerous species, including emblematic fish species such as walleye (*Sander vitreus*), yellow perch (*Perca flavescens*), northern pike (*Esox lucius*), and lake whitefish (*Coregonus clupeaformis*). These aquatic ecosystems support boreal and temperate communities shaped by diverse landscapes, such as the Canadian clay belt, eskers, and the Canadian Shield, and are influenced by strong land–water interactions (Grosbois et al., 2023).

**Figure 1.**
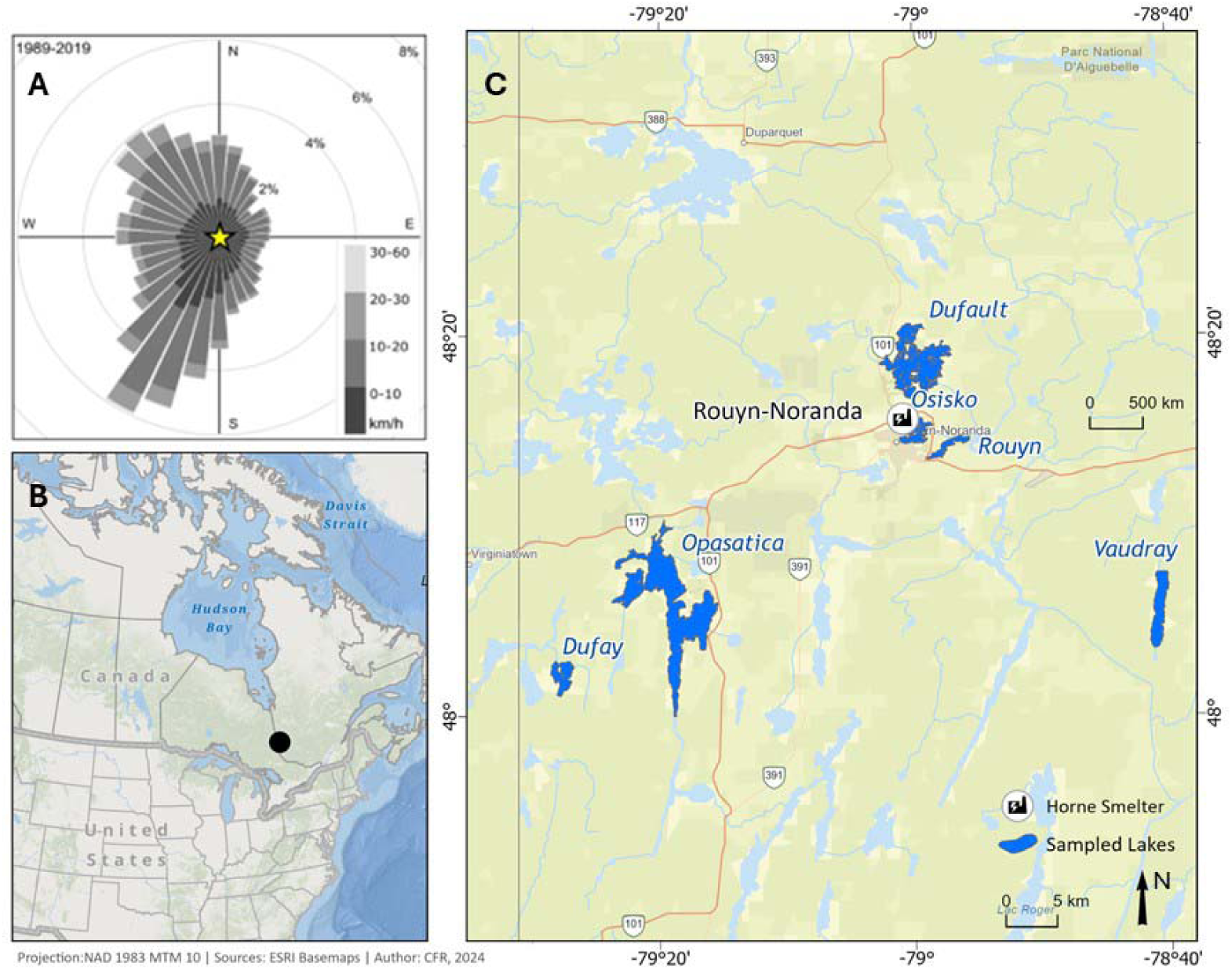
A - Wind rose estimated from 1989 to 2019, provided by the Meteorological Service of Canada, climate ID 7086716 et 7086720 (ECCC, 2019); B - Study area in the province of Québec, Canada; C - Study lakes (n = 6) in the Rouyn-Noranda region, also showing the localization of the Horne smelter.

Six lakes were selected (Fig. 1) along a gradient of contamination previously documented for the most studied TEs such as Cu, Cd, and Zn (Levesque et al., 2002; Giguère et al., 2004; Kraemer et al., 2008; Gauthier et al., 2009). These studies highlight spatial variations in metal deposition influenced by proximity to historical mining and smelting activities, such as those associated with the HS. Given these factors, three lakes close to the HS (Osisko (N1), Rouyn (N2), and Dufault (N3)) were selected due to their documented exposure to both direct discharges and diffuse contamination related to past industrial operations. In addition to metal contamination, N1 has been historically impacted by industrial and urban discharges, having once been used as a waste disposal site. In 1972, three dikes were constructed, dividing the lake into three sections and limiting water exchange between the newly formed basins. While the southern basin is now isolated from direct smelter discharges, it remains exposed to atmospheric deposition. In addition, the northern and central sections are still affected, receiving acidic water either from the smelter or from the municipal sewers of Rouyn-Noranda (Darricau et al., 2021). The latter is hydrologically connected to N2 through a small stream, extending the potential area of impact. N3, located north of the smelter, has also been exposed to both direct and atmospheric contamination despite being the drinking water source for the citizens of Rouyn-Noranda. In contrast, to assess contamination beyond the immediate industrial area, Vaudray Lake (I1), situated approximately 30 km downwind from the HS, was selected as an intermediate-distance site to investigate the potential for long-range atmospheric transport of metals. Finally, Opasatica (F1) and Dufay (F2), located farther from the smelter (30 km and 41 km upwind, respectively), were selected as sites documented in previous studies as being less impacted by mining activities. The physical and chemical characteristics of the lakes studied are summarized in Table 1.

### 2.2. Sample collection

In each lake, water, sediment and fish were sampled during the autumn of 2019, 2022 and 2023 (Supplementary information, SI; Table S1). Water samples were retrieved from the epilimnion (0.5 m below the surface) at three distinct locations of the lake using a Ruttner water sampler and a bucket, previously cleaned with nitric acid (HNO_3_; omni- trace Grade) at 15% (v/v) and rinsed seven times with ultrapure water (Milli-Q system; >18 MΩ · cm) to avoid unintentional TE contamination. Water samples (n=3/lake) for TE analyses were collected using metal-free 15 mL centrifuge tubes (VWR® International, Mississauga, ON) and then acidified with ultrapure HNO_3_ (2% v/v, Optima Grade). All samples were kept refrigerated at 4 °C until analyses. Water-column profiles of temperature (°C) and pH were measured using a multiparameter profiler (RBR Concerto, Ottawa, ON). Further details on water sampling protocols and analyses for other parameters, such as total nitrogen (TN), total phosphorus (TP), and other physicochemical variables, are provided in Table S2. Top sediments (2-3 cm) were sampled at each sampling site with an Ekman grab (15 × 15 × 15 cm) and placed in Whirl-pak plastic bags and frozen at −20 °C until analyses.

Fish individuals were collected in early June 2019 for F1 and N2, and in mid-September 2022 in F2, I1, N3, and N1. Two species, yellow perch and walleye were collected in the six studied lakes using gillnets on a single sample date per lake. The nets used were composed of eight panels (7.6 m long and 1.8 m height) disposed in increasing mesh order (25, 38, 51, 64, 76, 102, 127, and 152 mm). They were placed at depth ranging from 2–5 m and were left in the water for 18–24 h, always including the night. The number of nets used in each lake is determined with the lake area. Fish capture was done by our research team at the Université du Québec en Abitibi-Témiscamingue (UQAT) or in collaboration with the *Ministère de l’Environnement, de la Lutte contre les changements climatiques, de la Faune et des Parcs* (MELCCFP) (2022) and UQAM (2019).

For each 10 randomly selected fish per species (yellow perch and walleye), fork length and individual weight were recorded on the lake shore. Individuals were then stored at – 20 °C in resealable plastic bags until dissection in the lab. The liver and a dorsal muscle sample were taken from each fish with acid-cleaned forceps and scalpel and stored at −20 ° in plastic bags for TE analyses. Liver and muscle were selected as target organs due to their importance in detoxification processes and as a major site for energy metabolism, respectively. Additionally, muscle tissue is directly relevant to human consumption. We followed the protocol authorized by the animal protection committee (CPA) of UQAT (2022-03-31a) during the handling and euthanasia of net-caught fish and obtained a permit (2022-07-19-066-08-SP) from the MELCCFP. The fish morphological characteristics (e.g., sex, weight, fork length, condition factor) are given in Table S3.

### 2.3. Trace element analyses

The TEs, including TCEs, assessed in this study covered V, Cr, Fe, Co, Ni, Cu, Zn, As, Se, Sr, Mo, Ag, Cd, Sb, W, Ru, Pd, Pt, Tl, Pb, U, Ti, Y, La, Ce, Pr, Nd, Sm, Eu, Gd, Tb, Dy, Ho, Er, Tm, Yb, and Lu. All TEs levels in water, sediments and fish tissues were measured by a triple quadrupole inductively coupled plasma mass spectrometry (QQQ- ICP-MS, Agilent 8900, Santa Clara, CA) by the Laboratoire d’Analyses Environnementales (LAE) at UQAM. Detection limits and frequency detection are reported in Table S4.

Sediment samples were freeze-dried and homogenized before being analyzed. From 30 to 100 mg of dried sample was digested in tetrafluoroethylene vials with 6.4 mL of HNO_3_ (70%) and 1.6 mL of HCl (36%) using a single microwave digestion platform (Multiwave 5000, Rotor 24 HVT, Graz, AT). The initial cycle consisted of a 10-minute ramp-up period to 200 °C, followed by a 30-minute heating phase and a final 10-minute cooldown. The whole digestate (8 mL) was transferred to a 50-mL polypropylene trace metal-free centrifuge tube (VWR®) and the volume was completed to 40 mL with Milli-Q water (Lafrenière et al., 2025). Samples were diluted (1/10) with ultrapure water into trace metal-free tubes before analysis to obtain final HNO_3_ and HCl concentrations of 2% and 0.4 % (v/v), respectively.

Fish tissue (muscle and liver) samples were freeze-dried and homogenized before being analyzed for TEs. From 1 to 20 mg of dried weight (d.w.) was digested in a 15-mL polypropylene trace metal-free centrifuge tube (VWR®) with 500 µL of HNO_3_ (70%, Optima grade) for 24 hours. Samples were heated for 6 hours at 65 °C. Once cooled, 180 µL of grade hydrogen peroxide (30% H_2_O_2_, Optima grade) was added afterward for overnight digestion at room temperature. The volume was completed to 10 mL with Milli-Q water (Rosabal et al., 2015). Samples were diluted (1/2.5) with ultrapure water into trace metal-free tubes before analysis to obtain final HNO_3_ and H_2_O_2_ concentrations of 2% and 0.75 % (v/v), respectively.

For quality control, six blanks were included for each matrix: ultrapure water for water samples, and empty vials processed identically to the digestion procedure for sediment and fish tissues. Mean metal concentrations in blanks were subtracted from samples where the blanks were higher than the method detection limit (MDL).

The Certificated Reference Materials (CRM) MESS-4 (marine sediment, National Research Council of Canada, Ottawa, ON), TILL-3 (Geochemical Soil and Till Reference Materials, Certified Reference Materials Project, Ottawa, ON), BCR-667 (estuarine sediment, European Commission, Joint Research Centre, Geel, BE), and JSd-2 (stream sediment, Geological Survey of Japan, Tsukuba, JP) were used for sediment samples; while BCR^®^-668 (mussel tissue, European Commission, Joint Research Centre, Geel, BE), DOLT-5 5 (dogfish liver, National Research Council of Canada, Ottawa, ON), and 1517B (bovine liver, National Institute of Standards & Technology, Gaithersburg, MD), were used for fish tissues. Subsequently, the CRM were subjected to the same digestion method. Acceptable recovery percentages ranged from 81% to 126% (Supplementary information, SI; Table S4). Trace element concentrations are reported in µg per gram (dw) for sediments, in nmol per gram (d.w.) in biota, and in nmol per liter in water.

### 2.4. Data analysis

All data are represented with mean ± standard deviation (sd). Condition factors were calculated as (WL^-3^) × 100, where W is the fish body weight (grams) and L is the fish fork length (centimetres). Because the assumption of homogeneity of variance was not met for the raw data (Levene’s test), data transformations were attempted (e.g., log) to assess whether variance homogeneity could be achieved. However, some elements remained non-normally distributed even after transformation. Therefore, a non-parametric approach was adopted for all data analyses. Differences in hepatic and muscle TE concentrations among fish, as well as sediment and water TE concentrations among sampled lakes, were tested using the Kruskal-Wallis test. When significant differences were detected (*p* < 0.05), pairwise comparisons were conducted using Dunn’s test with Bonferroni correction. Compact letter display (CLD) groupings were derived from the post-hoc results to visualize significant groupings. When the concentration of a given element was below the MDL a multiplicative random lognormal replacement method was used (Palarea-Albaladejo et al., 2014). Wilcoxon rank sum tests were used to assess differences in TE concentrations between fish species. Statistical significance was considered at p < 0.05. The linear models were used to test the significance (p < 0.05) of the effect of proximity to the HS. These models included the TE concentrations as dependent variables and distance to the HS (in km) as the sole independent variable. Principal component analysis (PCA) was employed to determine and interpretate the main sources of TEs and further assess the comprehensive pollution. All statistical analyses and calculations were performed using R version 4.2.2 (R Core Team, 2016), with the packages ggplot2, ggpubr, dplyr, car, rcompanion, and FSA.

To assess the lakes environmental quality, we compared TEs concentrations in water and sediments to environmental quality guidelines. Specifically, we used the Canadian Water Quality Guidelines (CWQGs) for the protection of aquatic life where available (CCME, 2007). The lack of a Canadian guideline for REEs regarding water and sediment quality has led us to use the criteria proposed by the National Institute of Health and the Environment in the Netherland (Sneller et al., 2000). Similarly, Canadian Sediment Quality Guidelines (CSQGs) and Sediment Maximum Permissible Concentrations (SMPCs) were used to evaluate sediment contamination, while for several TCEs, guideline values remain unavailable.

To evaluate potential ecological and health implications, we measured TE concentrations in the muscle tissue of walleye and yellow perch collected from the six studied lakes. Guidelines for fish consumption (FAO/WHO, 2005) were used to assess the potential risks to human health; however, these reference values were only available for Cu, As, Zn, Cd, and Pb. Because these guidelines express concentration values in wet weight (ww), TEs concentrations in fish tissue were converted from dw to ww assuming an average water content of 80% (Begun et al., 2013).

## 3. Results

### 3.1. Contamination patterns for historical and emerging contaminants

#### 3.1.1. Trace element concentrations in lake water and sediment

Most TEs, including TCEs and REEs, were quantified in water and sediment samples (detection frequencies higher than 88%) except for Ru, Pt, W, and Pd, which were below detection limits (Supplementary information, SI; Table S4). Water and sediment concentrations of TEs varied across lakes with maximum-to-minimum ratios ranging from 6 (Sr) to 386 (Ce) in water, and from 1 (Sr, Nd, Ce) to 268 (Ag) in sediments (Table S5). Notably, elements such as La, Ce, Nd, Pr, Tl, and Ti showed the highest spatial gradients in water, while Ag, Sb, Tl, Cd, and Cu exhibited the greatest variation in sediments. In general, the three lakes closest to the HS (N1, N2, N3) were consistently more contaminated in water than both F1 and F2, the farthest ones to the contamination source (Fig. 2; Tables S6-S7). Mean concentrations of Cu, Zn, As, Se, Cd, and Pb were lower in water from N1 compared to N2 and N3, but higher than those found in water from F1 or F2. For instance, Cd concentration in N1 was 6.5 times lower than in N3, but 4.3 times higher than F1. While the highest concentrations for Cu, Zn, As, Se, and Cd were found in N3, the lowest levels were measured in F1 (Zn, As, Se, Cd) and F2 (Cu). Similar to the water samples, the same three lakes closest to the smelter consistently exhibited the highest sediment concentration, in contrast to the two more distant lakes (F1 and F2) (Fig. 3; Tables S8-S9). Lake N1 showed the highest sediment concentrations for Cu, Zn, As, Se, and Cd, whereas F1 reported the lowest values. For instance, As sediment concentrations in F1 were about 79 times lower than in N1. Similarly, I1 and F2 lakes showed lower concentrations of the previously reported TEs compared to those located closer to the HS. For example, the average concentrations of Se in sediment from I1 and F2 lakes were approximately 15 to 134 times lower than those measured at the most contaminated sites. Similarly, Cu concentrations in lake I1 and F2 decreased by 96% and 99%, respectively, while Zn levels were reduced by 86% in I1 and 95% in F2 compared to lakes located close to the smelter.

**Figure 2.**
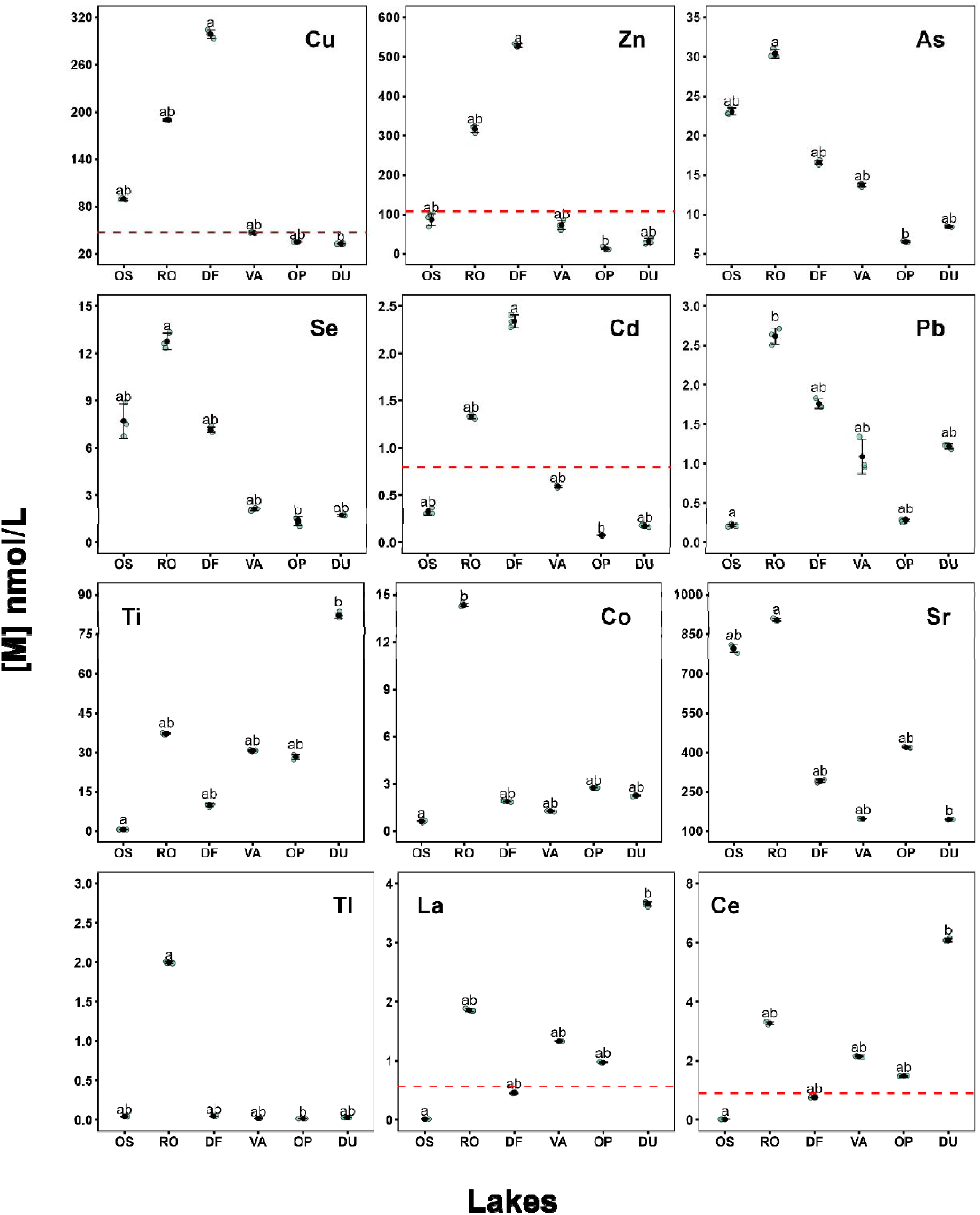
Water concentrations of Cu, Zn, As, Se, Cd, Pb, Ti, Co, Sr, Tl, La, and Ce (n = 3; nmol/L) were measured in six lakes categorized by proximity to the Horne Smelter (nearby: N1–N3; intermediate: I1; distant: F1–F2). Data are presented with error bars indicating the standard deviation. The red dotted line indicates the Canadian Water Quality Guidelines (CWQGs) for the protection of aquatic life from dissolved metals (not visible for As: 66.97 nmol/L; not available for Se) (CCME, 2007) or the background concentrations in water for REEs proposed by the National Institute of Health and the Environment in the Netherlands (Sneller et al., 2000). The background concentrations are not available for Ti, Co, Sr, and Tl. Different letters indicate significant differences between means (Kruskal-Wallis, followed by Dunn honestly post hoc test on ranks, *p* < 0.05).

**Figure 3.**
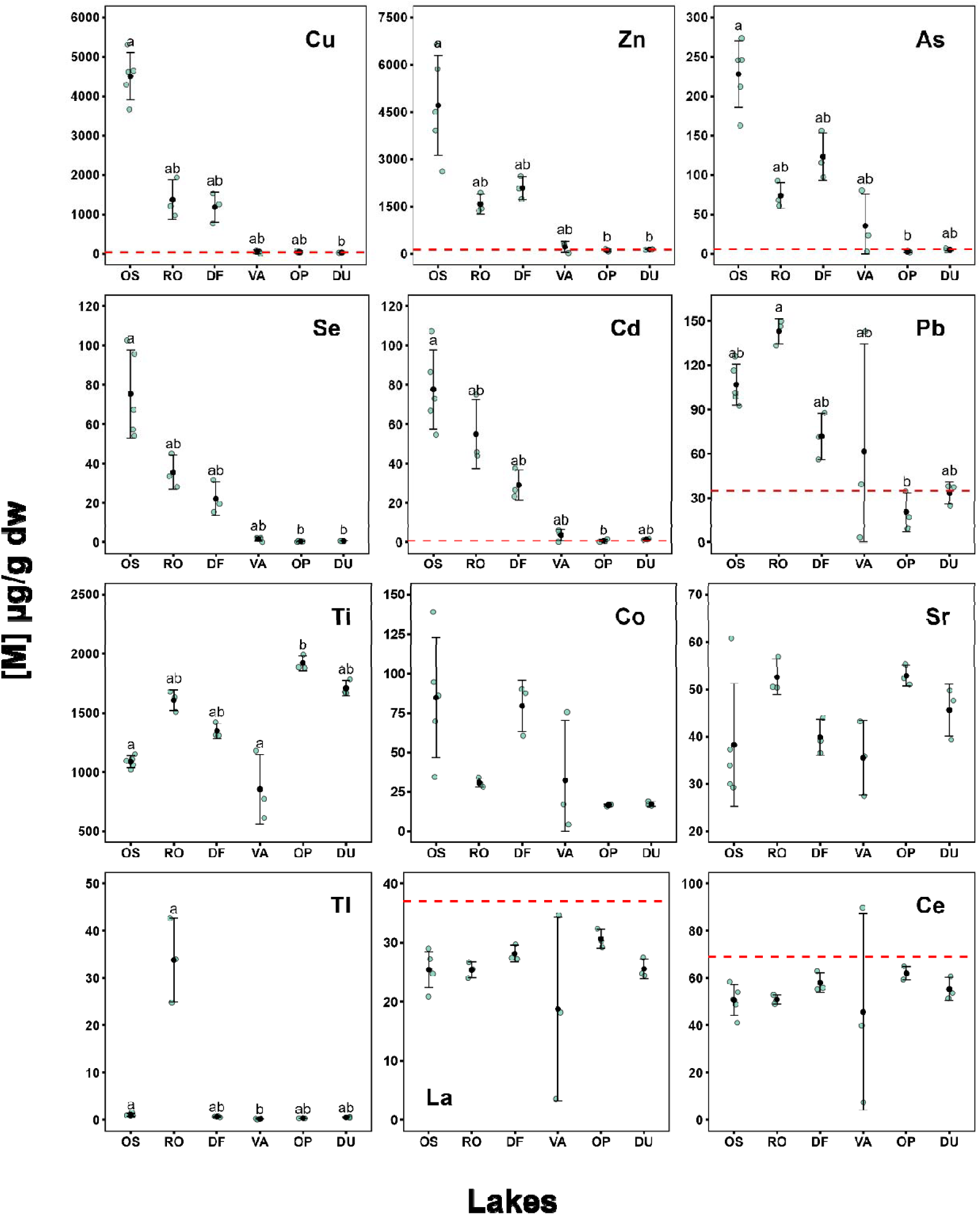
Sediment concentration (n=3-5; µg/g dry weight, dw) of Cu, Zn, As, Se, Cd, Pb, Ti, Co, Sr, Tl, La, and Ce were measured in six lakes categorized by proximity to the Horne Smelter (nearby: N1–N3; intermediate: I1; distant: F1–F2). Data are presented with error bars indicating the standard deviation. The red dotted line indicates the Canadian Sediment Quality Guidelines (CSQGs) for the protection of aquatic life (not available for Se) (CCME, 2007) or the background concentrations in sediment for REEs proposed by the National Institute of Health and the Environment in the Netherlands (Sneller et al., 2000). The background concentrations are not available for Ti, Co, Sr, and Tl. Different letters indicate significant differences between means (Kruskal-Wallis, followed by Dunn honestly post hoc test on ranks, *p* < 0.05).

For TCEs, we observed spatial variation in the concentrations of Ti and Tl in both water and sediment, while Co, Sr, La, and Ce varied significantly only in water (Figs. 2– 3). Water concentrations of Co, Sr, and Tl were elevated in N2 compared to the other lakes (e.g., up to 6.5 times higher for Co and Sr than those measured in F2). An opposite trend was observed for Ti and the two REEs, La and Ce, which showed the highest concentrations in F2. For instance, La concentrations in F2 reached 3.6 nmol/L, while only 0.01 nmol/L was detected in N1, representing a 360-fold difference (Fig. 2). In sediments, no spatial differences among lakes were detected for Co, Sr, La, and Ce. However, contrasting patterns were still observed for certain elements. Tl concentrations in N2 sediments were approximately 103 times higher than in F1 and 77 times higher than in F2, while Ti displayed the highest levels in F1 compared to N1 (approximately 72% higher) (Fig. 3).

Proximity to the HS clearly influenced the concentration of several TEs in both water and sediment (Table S10). In water, we found negative correlations between the distance from the HS and concentrations of Cu, Zn, Cd, As, and Se (*p* < 0.01), as TEs levels decreased with greater distance from the smelter. Same trend was observed for the TCE Sr (*p* < 0.001; Table S10). On the contrary, Pb concentrations were independent of distance, as the TCEs Co and Tl (*p* > 0.05), implying that their distribution is likely influenced by other environmental drivers. Surprisingly, we observed the highest concentrations for the REEs La, Ce, and Ti farther from the smelter (*p* < 0.01). In sediments, Cu concentrations declined by approximately 91 µg/g dw per kilometer away from Rouyn-Noranda (*p* < 0.001; Table S10). A similar spatial pattern was observed for Cd, whose levels were approximately 61 times lower in the farthest lake (F2) compared to the closest (N1). We also found negative correlations with distance for Zn, As, Se, Pb, and Co (*p* < 0.01). Conversely, elements such as Ti, Sr, Tl, La, and Ce remained invariable across the gradient (*p* > 0.05).

#### 3.1.2. Trace element concentrations in fish liver

Liver of walleye and yellow perch contained TEs such as Cu, As, Zn, Cd, Fe, Se, and Pb, while Cr, Ni, Ag, Mo, and U were not detected in most samples (MDL frequencies below 88%). Among TCEs and REEs, Sb, Pd, W and Gd, Nd were also absent in fish liver (Table S4).

Consistent to the patterns observed in water and sediment, TEs associated with historical mining activities were found at higher concentrations in both fish species collected in lakes close to the HS compared to those from more distant sites (Figs. 4-5; Tables S11-S12). In N1, the mean Cu levels in walleye fish liver were approximately 3.3 times higher than in F1, 3.1 times higher than in I1, and 2.4 times higher than in F2. Selenium levels showed even more pronounced differences, being 6.3 times higher in fish from N1 than those in individuals from F1, 4.5 times higher than in I1, and 3 times higher than in F2. In addition, for Zn, As, and Cd, mean levels measured in N1 were consistently the highest concentrations reported among lakes (e.g., N1: 5 µg/g dw; N2: 1 µg/g dw). In yellow perch liver, concentrations of Cu, Zn, Se and Cd were highest at site N1 compared to all other sampled locations. For instance, Cd concentrations in N1 were approximately 364% higher than in F1 and 359% higher than in F2. Similarly, Se concentrations in walleye fish livers from N1 were 222% higher than in F1, 145% higher than in F2, and 131% higher than in I1.

**Figure 4.**
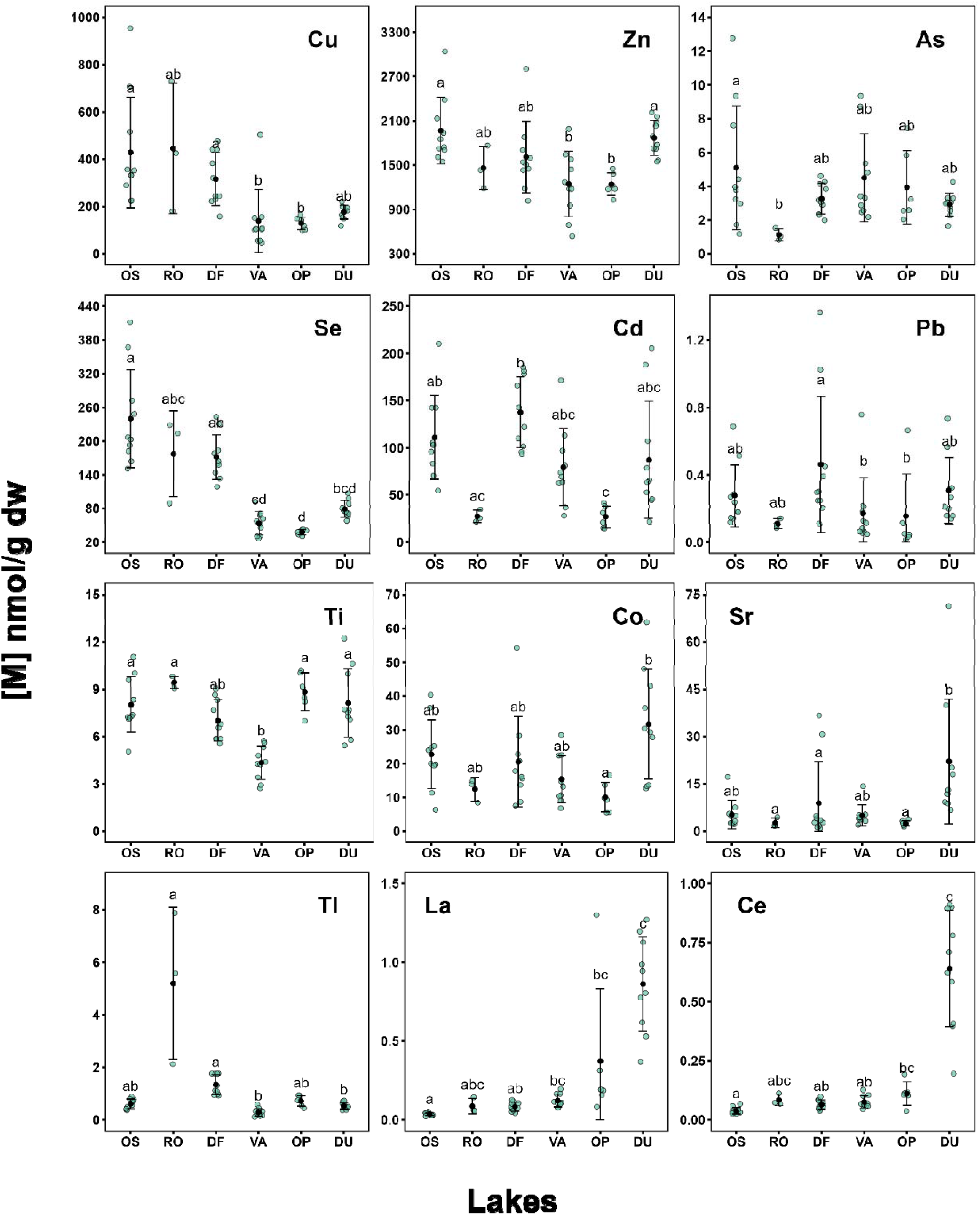
Concentration of Cu, Zn, As, Se, Cd, Pb, Ti, Co, Sr, Tl, La, and Ce in walleye’s liver (n=3-10; nmol/g dry weight, dw) were measured in six lakes categorized by proximity to the Horne Smelter (nearby: N1–N3; intermediate: I1; distant: F1–F2). Data are presented with error bars indicating the standard deviation. Different letters indicate significant differences between means (Kruskal-Wallis, followed by Dunn honestly post hoc test on ranks, *p* < 0.05).

**Figure 5.**
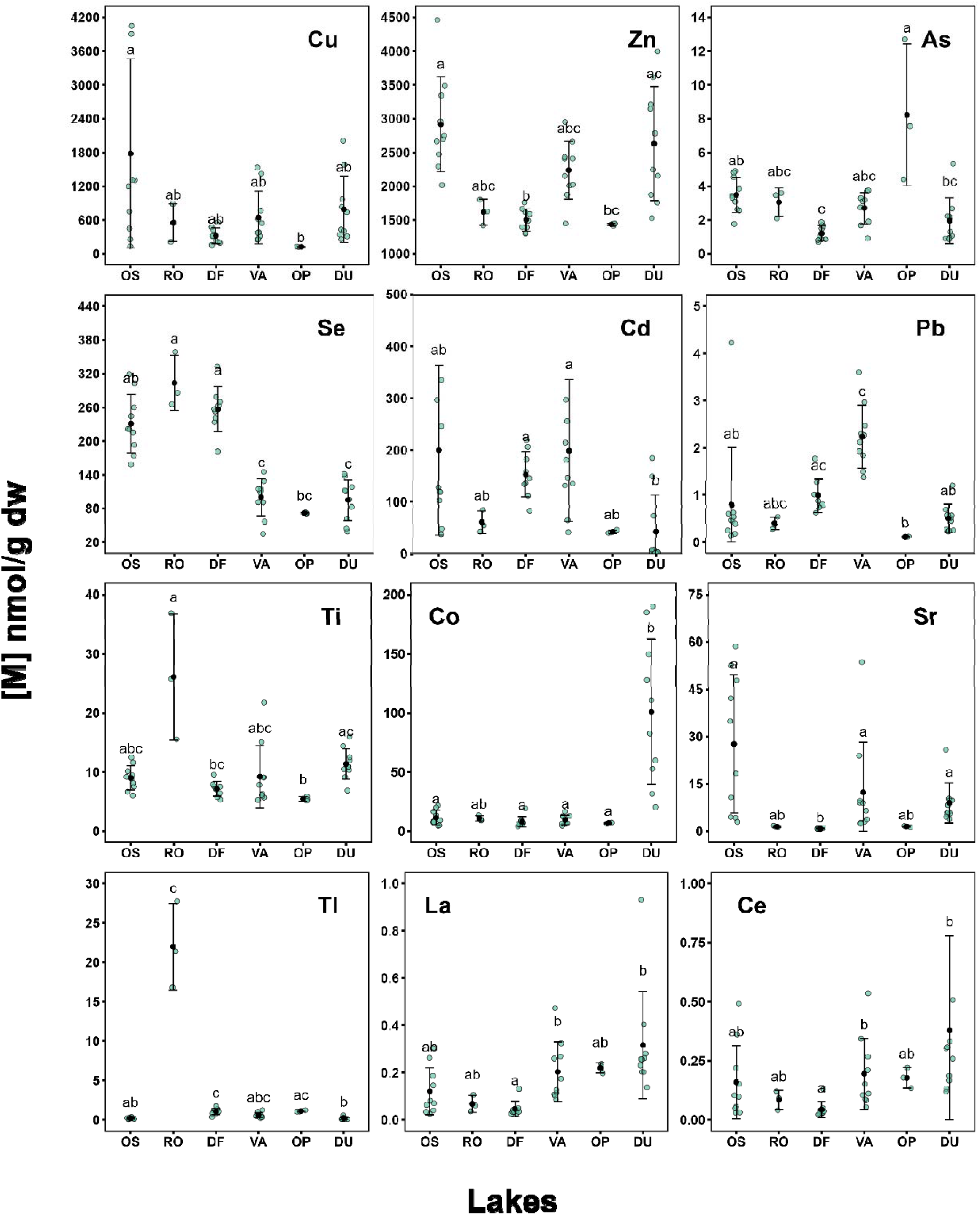
Concentration of Cu, Zn, As, Se, Cd, Pb, Ti, Co, Sr, Tl, La, and Ce in yellow perch’s liver (n=3-10; nmol/g dry weight, dw) were measured in six lakes categorized by proximity to the Horne Smelter (nearby: N1–N3; intermediate: I1; distant: F1–F2). Data are presented with error bars indicating the standard deviation. Different letters indicate significant differences between means (Kruskal-Wallis, followed by Dunn honestly post

In contrast to the more frequently studied TEs, TCEs exhibited distinct spatial distribution patterns in fish across lakes. While Ti and Tl showed higher concentrations in both fish species from N2, other elements such as Co, La, Ce, and, in the case of walleye, Sr, displayed elevated levels in fish from lakes farther from the HS (Figs. 4-5; Tables S11-S12). For instance, Ti concentrations in walleye liver collected from N2 were modestly higher (about 12.5%) than in N1 and F2, while Tl concentrations were markedly elevated–approximately 843%, and 733% greater than in N1 and F2, respectively. On the contrary, La and Ce mean levels in walleye fish liver were 11 and 8 times higher in F2 compared to N2, respectively (Fig. 4). Similarly, Sr levels in fish collected from F2 were 4.4 higher than N1, and 8.2 higher than N2. In yellow perch liver, Ti mean concentrations in N2 were approximately 2.4 and 1.6 times higher than those measured in N1 and F2, respectively. For Tl, concentrations in N2 were substantially amplified–approximately 122 times greater than in N1 and F2 lakes. Similar to walleye liver, yellow perch La and Ce concentrations in F2 were higher than in N1, respectively around 160% and 138%. Conversely, Sr levels in N1 were about 200% higher than in F2, highlighting a contrasting pattern between the two species.

For both walleye and yellow perch, TEs concentrations in liver tissues showed species-specific relationships with distance from the HS (Table S13). Regarding walleye liver, negative correlation was observed between distance from the smelter and Cu, Se, and Cd concentrations (*p* < 0.05), as metals levels diminished as the distance expanded. In contrast, concentrations of Zn, As, and Pb were not significantly related to distance (*p* > 0.05). In yellow perch liver, Se levels showed a significant negative correlation with distance from the HS (*p* < 0.001). Other elements such as Cu, Zn, As, Cd, and Pb were invariant regardless the distance (*p* > 0.05). For TCEs, spatial trends also varied between species (Table S13): walleye liver Tl concentrations declined with increasing distance (*p* < 0.05), whereas La and Ce showed higher concentrations in more distant lakes (*p* < 0.001). No trends were observed for Ti and Co (*p* > 0.05). Similarly, in yellow perch, La and Ce concentrations also increased with distance (*p* < 0.001), as did Co, whereas Ti, Sr, and Tl were consistent at all distances (*p* > 0.05).

When comparing liver TEs concentrations between the two species across all six lakes, yellow perch showed higher levels than walleye. For instance, yellow perch showed higher liver concentrations of Cu, Pb, Zn (all *p* < 0.001), Se (*p* < 0.01), and As, Co, Tl, Ti (all *p* < 0.05) compared to walleye. In addition to interspecific differences, individual traits influenced TE concentrations. Liver Se concentrations increased with length in both fish species (*p* < 0.05; Fig. S1), while Cu and Se were elevated with both length and weight only in walleye (*p* < 0.01). For other TEs such as As, Zn, and Pb as well as TCEs like Ti, Co, Sr, La, and Ce no significant relationship with fish size (length or weight) was observed in either species (*p* > 0.05).

### 3.2. Environmental quality assessment

#### 3.2.1. Water and sediment quality

Of the 6 lakes included in our analysis, 4 of them exceeded the CWQGs for the protection of aquatic life from dissolved metals in lake water, for at least one of Cu, Zn, or Cd. Water Cu concentrations exceeded the CWQGs (47 nmol/L) (Fig. 2; Table S6) in all lakes except for F1 and F2 lakes. Similarly, Zn and Cd concentrations in water lake exceeded the CWQGs (107 and 0.8 nmol/L, respectively) in N2 (Zn: 322 nmol/L; Cd: 1.3 nmol/L) and N3 lakes (Zn: 527 nmol/L; Cd: 2.3 nmol/L). In contrast, As concentrations remained below the CWQGs of 66 nmol/L across all lakes. The CWQGs for Se, Pb, and other elements such us Fe and Ni are not currently available in Canada (Table S6). The REEs La and Ce exceeded the WMPCs in all lakes except for N1 and N3 lakes, where concentrations were below 0.57 and 0.92 nmol/L, respectively (Fig. 2). However, no international water quality guidelines are currently available for other TCEs such as Ti, Co, Sr, and Tl (Table S7).

Sediment concentrations of Cu, Zn, As, Pb, and Cd surpassed the CSQGs for the protection of aquatic life in all sampled lakes (e.g., N1 exceeded 12,733 times Cd’s guidelines), apart from of Zn and Pb in F1 (with an average value of Zn about 10% lower than the guideline; Table S8). The median sediment concentrations of the REE La and Ce were below the SMPCs in all the lakes. No international sediment quality guidelines are currently available for Tl and other TCEs such Co, Sr, and Ti (Table S9).

### 3.3. Human health risk assessment

#### 3.3.1. Trace element concentrations in fish muscle

Trace elements such as Cu, Zn, and Se were present across muscle samples of both walleye and yellow perch, whereas others, including Cr, Ni, Ag, Mo, and U, were frequently below detection limits (detection frequencies <88%). Similarly, among TCEs and REEs, the elements Sb, Pd, W, Gd, and Nd were undetected in muscle (Table S4).

For both fish species, permissible limits for the TEs As, Zn, and Pb were exceeded in all lakes for at least one of these elements, with some lakes showing multiple exceedances. In walleye muscle tissue, elevated mean concentrations of several TEs were observed across both close and distant lakes, including Cu (7 and 6 mg/kg ww in N1 and F2, respectively), As (2.4 and 0.9 mg/kg ww in I1 and F1, respectively) Zn (107 and 99 mg/kg ww in N1 and F2, respectively), Cd (0.09 mg/kg ww in N3), Se (42, 27, and 29 mg/kg ww in N1, N2, and N3, respectively), and Pb (0.5 mg/kg ww in N1) (Table 2). Mean Zn concentrations in N1 and N2 exceeded the permissible limit of 100 mg/kg ww, whereas mean values in the other lakes remained below this limit. Nevertheless, individual exceedances were recorded in the farthest lakes as well, with a maximum Zn value of 134 mg/kg ww observed in F2. For Pb, the mean concentration in N1 matched the FAO/WHO guideline value of 0.5 mg/kg ww, while all other lakes exhibited lower mean levels. However, as for Zn, some fish from other lakes also exceeded this threshold, with a maximum Pb concentration of 0.8 mg/kg ww recorded in N3 and 0.5 mg/kg ww in F2. In the case of As, the mean concentration in walleye liver from I1 was 2.4 mg/kg ww, more than twice the recommended limit of 1 mg/kg ww. In N2, individual concentrations also surpassed the limit, with maximum values reaching 1.3 mg/kg ww. No international safety thresholds are currently established for Se in fish tissue, complicating the evaluation of potential health risks associated with this element. In contrast, Cd and Cu concentrations in walleye liver remain consistently below the FAO/WHO limits across all lakes. For instance, the Cu concentration in N1 (7 mg/kg ww) was approximately 77% below the recommended limit of 30 mg/kg ww. In yellow perch liver, Zn means concentrations exceeded the FAO/WHO permitted limit of 100 mg/kg ww in all lakes. For this element, the highest concentration was observed in N1 (302 mg/kg ww), representing a 202% exceedance of the limit. For Pb, all lakes except N2 and F1 exhibited mean concentrations above the guideline value of 0.5 mg/kg ww. The highest mean Pb concentration was recorded in F2 (3.7 mg/kg ww), over seven times the permitted limit. As in walleye, the highest mean As concentration in yellow perch liver was observed in I1 (1.0 mg/kg ww), exactly matching the limit value. In contrast, the mean concentrations of Cd and Cu remained well below recommended thresholds across all lakes. However, some individuals from I1 and F2 showed Cu concentrations exceeding the limit.

For TCEs and REEs, as observed for TEs, elevated concentrations were found in both close and distant lakes. However, comparisons with FAO/WHO permitted limits remain limited due to the lack of established guidelines for these elements. In walleye, mean concentrations of Ti and Co were highest in N1 (Ti: 2.8 mg/kg ww; Co: 0.6 mg/kg ww), while Sr peaked in N2 (6 mg/kg ww), and Tl in F2 (2.7 mg/kg ww). Yellow perch showed elevated Ti in F1 (5.3 mg/kg ww), highest Co and Tl levels in N2 (Co: 0.3 mg/kg ww; Tl: 3.8 mg/kg ww), and Sr in I1 (33 mg/kg ww). For RREs, in both species, the highest mean concentrations of La and Ce were found in F2 (e.g., La: 0.4 mg/kg ww in walleye, 0.07 mg/kg ww in yellow perch).

TEs concentrations in muscle tissue differed significantly between the two fish species. Yellow perch generally exhibited higher levels than walleye for most elements, including Cu, As (*p* < 0.001), Cd (*p* < 0.0001), Zn (*p* < 0.0001), Pb (*p* < 0.0001), Se (*p* < 0.01), Co (*p* < 0.0001), Tl (*p* < 0.001), Ti (*p* < 0.01), Sr (*p* < 0.0001), and Ce (p < 0.05).

No significant difference was observed for La (*p* > 0.05). In addition to interspecific differences, individual traits influenced TE concentrations. Muscle Se concentrations increased with length and weight in both fish species (*p* < 0.05; Fig. S2), while Ti and Co concentrations increased with length only in walleye (*p* < 0.01). For other TEs such as As, Zn, and Pb as well as TCEs like Ti, Co, Sr, La, and Ce no significant relationship with fish size (length or weight) was observed in either species (*p* > 0.05).

### 3.4. Spatial clustering of lakes based on trace elements

The PCA consistently revealed a clear spatial segregation of lakes according to their proximity to the HS across all analyzed matrices (water, sediment, and liver tissue of both walleye and yellow perch) (Fig. 6-7). In each case, lakes located closer to the contamination source (N1, N2, and N3) were distinguished from more distant lakes (I1, F1, and F2).

**Figure 6.**
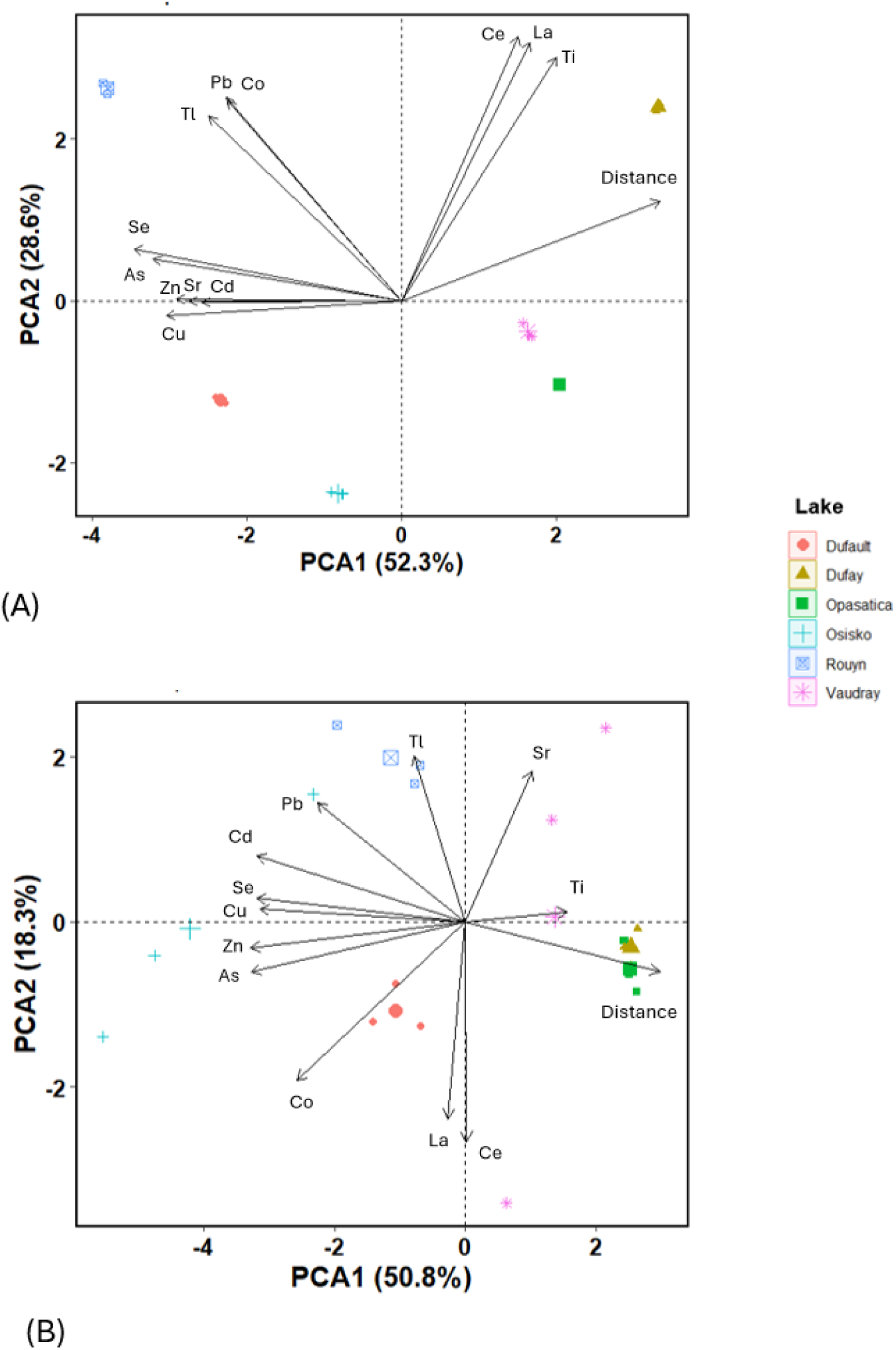
Principal component analysis of trace elements in surface water (A) and sediments (B) from the six studied lakes. PC1 and PC2 accounted for 52.3% and 28.6% of the variance in water, and for 50.8% and 18.3% in sediments.

**Figure 7.**
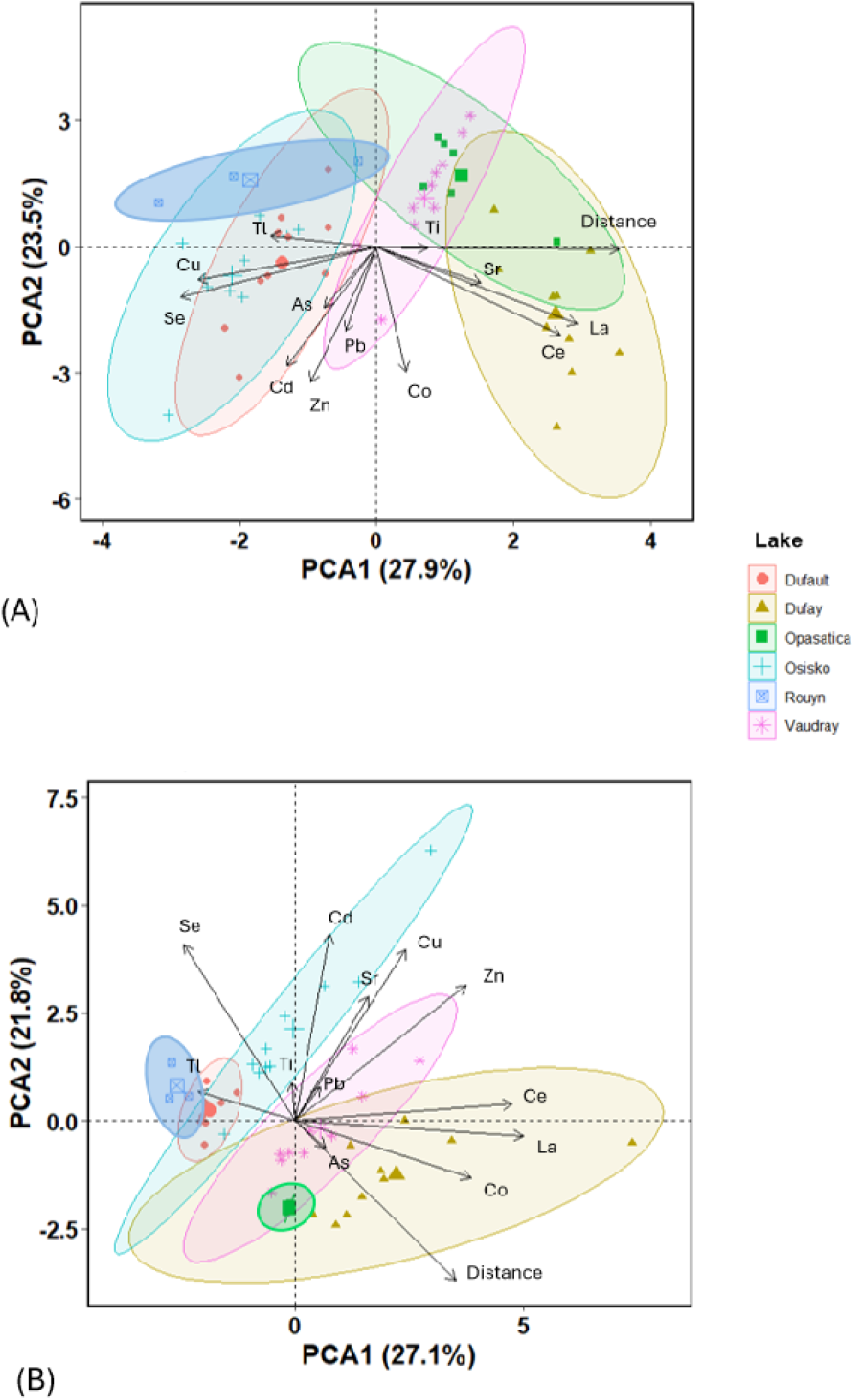
Principle components of trace elements in the walleye (A) and in yellow perch liver (B) from the six studied lakes. PC1 and PC2 accounted for 27.9% and 23.5% of the variance in walleye, and for 27.1% and 21.8% in yellow perch.

In water, Cu, Zn, As, Se, Cd, and Pb were the main contributors to PC1, all showing negative correlations with this axis (e.g., Se, r = -0.36; see Fig. S3A in supplementary material). However, Pb exhibited a lower negative loading on PC1 (-0.23) than elements such as Se (-0.36) or As (-0.33) and a higher positive loading on PC2 (Pb: 0.35; Se: 0.09). In regard to TCEs, Co and Tl were positively correlated in water (0.99; Fig. S4A) and showed a similar pattern of variation as the previously mentioned TEs, being positively associated with PC2 (e.g., Co, r = 0.35) and negatively with PC1 (e.g., Co, r = - 0.23). Sr was positively correlated with elements such as Se, Cu, As, Cd, and Zn ((e.g., Sr–Se: r = 0.82; Sr–As: r = 0.82; Fig. S4A) and similarly showed a negative correlation with PC1 (-0.27). Ce, La, and Ti, which were strongly correlated in water (r = 0.99), contributed mainly to PC2 (e.g., La: 0.45 on PC2; La: 0.17 on PC1 Fig. S3A) and were positively associated with distance from the HS (e.g., La–distance: r = 0.69; Fig. S4A).

In sediment, ss for the water matrix, Cu, Zn, Cd, As, and Se were also the main contributors to PC1 and showed negative loadings on this axis (Zn: -0.37; Fig. S3B). These elements in sediment were positively correlated with each other (e.g., Cu–Zn: r = 0.95; Fig. S4B). On PC2, Cu, Se, and Cd showed positive loadings (e.g., Cd: 0.15) whereas Zn and As loaded negatively (e.g., As: –0.11). As in water, Pb was correlated with the other TEs (e.g., Pb–Se: r = 0.58), showing a negative loading on PC1 (–0.25) and a positive loading on PC2 (0.27). Similarly, Sr and Tl also loaded positively on PC2 (0.34 and 0.38, respectively). In contrast, Co appears lower in the same quadrant, with negative correlation for both PCs (loads for PC1 and PC2: -0.29 and -0.36, respectively). La and Ce showed a negative correlation with PC2 (Ce: -0.51; La: -0.45). Ti exhibited the highest positive correlation with distance from the HS (r = 0.25).

For walleye liver, as for the water and sediment matrices, Cu, Zn, Cd, As, and Se showed negative loadings on PC1 (Cu: -0.36; Figs. 7A-S3B). These elements in walleye liver were also positively correlated with each other (e.g., Cu–Zn: r = 0.76; Fig. S4C). PC1 separates lakes according to the distance from the source of contamination: closer lakes are positively associated with elements such as Cu, As, Se, Zn, Cd, and Pb, while farther lakes are associated with elements such as Sr, La, and Ce (distance-Ce: 0.65; Fig. S4C). The variable distance was positively correlated with PC1 (r = 0.49; Fig. S3C). PC2 separates samples based on high loadings of Tl and low (negative) loadings of Co (Fig. 7A).

As for yellow perch liver, Cu, Zn, Pb, and Cd showed positive loadings on both PC1 (e.g., Cu: 0.23) and PC2 (e.g., Cu: 0.42) (Figs. 7B-S3D). The main contributors to PC2 were Cd (0.46), Se (0.43), Cu (0.42), and Zn (0.33), elements typically associated with industrial pollution. Here, the distance variable had a moderate loading on PC1 (0.33) but a strong negative loading on PC2 (–0.39). Elements such as Se and Cd contributed more strongly to PC2 than to PC1 (e.g., Se: PC2 = 0.43 vs. PC1 = –0.23). PC1 presented positive loadings also for La (0.47), Ce (0.45), and Co (0.36), as well as a loading of 0.33 for the distance to the HS (Fig. S3D). Notably, the distance variable loaded negatively on PC2 (–0.39). Co, La, and Ce were positively correlated with distance (e.g., distance–Co: r = 0.54), whereas other TCEs such as Ti, Tl, and Sr showed negative correlations with this variable (Fig. S4D).

## 4. Discussion

### 4.1. Spatial distribution of trace element and probable sources

#### 4.1.1. Trace element in lake water and sediments

Our results evidenced a significant spatial differentiation in the distribution of historically reported TEs such as Cu, Zn, As, and Cd, as well as some emerging TCEs (i.e., Tl, Co, and Sr) but not for REEs, in water and sediments of lakes located close to the HS compared to more remote sampled lakes. The spatial gradient observed (N1, N2, N3 > I2 > F2 > F1) suggests a strong anthropogenic influence related to historical mining activities and recent atmospheric emissions from the copper smelter. When we compared our results to historically monitored TEs with studies from the early 2000s, we observed patterns in both water and sediment that are consistent with those previously reported in the Rouyn-Noranda region. In fact, water samples collected in 2001 and 2003 showed a similar pattern of TE concentrations (Giguère et al., 2005; Kraemer et al., 2006). In the same year, Couture and Pyle (2008) expanded this assessment by analyzing both water and sediment from a set of lakes that included one distant site (our lake F1), one intermediate lake, and two lakes closer to the smelter (N1 and N3). They also reported an increase in TEs concentrations with decreasing distance from the smelter. In a survey of lake sediments near the HS, TEs concentrations in surface sediments decreased with distance from the source along the path of the prevailing wind, underscoring the role of atmospheric deposition in shaping spatial patterns (Telmer et al., 2006). Similarly, Borgmann et al. (2004), working in the same area, reported Cu levels in the lake closest to the smelter (N3) to be up to 40 times higher than in lakes situated 22–51 km away. Other TEs, including Pb and Zn also showed substantial declines with increasing distance from the source (Borgmann et al., 2004). This differential distribution agrees with findings from other mining regions and industrial activities worldwide. For example, Simmatis et al. (2022) in Manitoba and Gashkina et al. (2015) in Russia reported marked enrichment of Cu, Zn, and Cd in lake sediments and water downstream of Cu smelters, with concentrations decreasing progressively with distance from the emission source. Likewise, Hadjipanagiotou et al., (2020) observed significant enrichment of Cu, Zn, Pb, and Cd in sediments of pit lakes and stream surrounding the abandoned copper mine in Agrokipia (Cyprus). In their study, TEs concentrations in stream waters showed a decreasing trend with increasing distance from the tailings dump, reaching background values within 1,500 meters.

In our study, the less-studied Tl, Co, and Sr have also exhibited similar spatial pattern to those historically monitored TEs suggesting that these elements, although less studied in the past, may follow contamination patterns similar to those of better-known metals. In particular, the elevated level of Sr at the closest sites, along with its strong correlation with commonly monitored metals in water (e.g., As), suggests a shared industrial origin. However, the low and spatially uniform concentrations of Sr in sediments suggest that this element remains more mobile in the water column and is less likely to accumulate in the benthic environment (Boyer et al., 2018). This characteristic may, in fact, reflect a high bioavailability and therefore pose an underestimated ecological risk for aquatic biota. HS’s increasing involvement in e-waste treatment in recent decades may have contributed to the introduction of new contaminants into lakes within the Rouyn-Noranda area (Dupont et al., 2025; Ponton et al., 2016), with yet little information available but potentially significant ecological implications. Recycling processes are known contributors to environmental TE contamination (LaCoste et al., 2001; Song and Li, 2014). Supporting this, Proulx et al. (2015) highlighted a strong spatial gradient in Tl concentrations across Rouyn-Noranda lakes, with levels in N2 exceeding regional background values by over 300 times, and more than 330 times those observed in N1. Similarly, Borgmann et al. (2004) reported that, already 14 years after the start of the new recycling activity from the HS, Tl concentrations in surface sediments of lakes located close to the HS were 2.8–6.8 times higher than in more distant lakes, reinforcing the influence of smelting emissions on Tl distribution. The frequent co-occurrence of Tl and Co with Pb suggests that these elements may be transported and deposited through similar pathways, such as atmospheric deposition of metal-laden fine particulates as well as via industrial effluents. Their presence across multiple environmental matrices underscores both the extent of anthropogenic influence and the growing importance of monitoring emerging contaminants. Beyond the Rouyn-Noranda area, studies in other metal-impacted regions have revealed similar, though sometimes contrasting, spatial patterns for TCEs. For instance, Bentley et al., (2023) documented a clear spatial variability in the Great Lakes basin waters, with Tl and Co concentrations decreasing downstream from major anthropogenic sources, indicating a strong contribution from urban and industrial point sources. However, Soroaga et al. (2022) reported that although Co and Tl were detected in lake sediments from an industrial area in northeastern Romania, they did not exhibit significant enrichment, suggesting a relatively low degree of contamination of the area. This discrepancy highlights how TCEs can respond differently to industrial inputs depending on their geochemical behavior, local environmental conditions, and the nature of emission sources.

In contrast, Ti and REEs (La and Ce) exhibited an opposite pattern in our study. Ti showed higher concentrations in the farthest lakes from the HS and lower concentrations in lakes closer to the contamination source, following the trend F2 > F1, I1, N3, N2 > N1, consistent in both water and sediments. For La and Ce, this trend was observed in water but not in sediments, where no significant variation among lakes was detected. This contradicts our hypothesis of an anthropogenic influence. Indeed, the co- variation between Ti, La, and Ce in water, distinct from those of other TEs, further suggests different environmental drivers such as rock composition, erosion, or runoff processes. Their distribution appears therefore to be more influenced by local lithology and soil characteristics rather than anthropogenic sources. Ricard-Henderson (2023) recently reported that REE distribution in water and *Chaoborus* larvae in lakes from two mining regions, Rouyn-Noranda and Sudbury (Ontario, Canada), mainly reflects natural geochemical processes, including processes such as oxidation, precipitation, and complexation, rather than anthropogenic activities. Similarly, Dupont et al. (2025) investigated the atmospheric deposition of both historically monitored and emerging metals, including REEs, in terrestrial ecosystems surrounding the HS in Rouyn- Noranda. Their results showed limited evidence of REE enrichment in biomonitors and passive air samplers, suggesting minimal impact from smelting emissions on REE contamination in nearby forested environments. Comparable results have been reported by Zubova et al. (2020) who documented REE concentrations up to 14 times higher than local background levels in Arctic lakes of Northwestern Russia. These concentration anomalies were largely attributed to underlying geology, particularly the presence of bedrock enriched in Ce and La. In addition, in the mining-impacted region of Bow Lake (Ontario), Dang et al. (2021) found extremely high La concentrations (up to 2200 µg/g dw), strongly correlated with uranium content and microbial processes such as methanotrophy. The co-precipitation of REEs with phosphorus minerals and their partial association with organic matter suggested complex biogeochemical mechanisms behind their retention in sediments. Together, these comparisons reinforce the interpretation that REE and Ti concentrations in our farther lakes primarily reflect natural geochemical conditions, such as runoff from REE-rich soils or erodible substrates, rather than industrial inputs. This distinction is also supported by the relatively low accumulation of these elements in sediments compared to their mobility in water, indicating weaker sorption or sedimentation tendencies. The contrasting behavior between water and sediment for elements like Sr, La, and Ce likely reflects their differing affinities for particulate matter, solubility, and chemical speciation, with implications for their ecological mobility and potential bioavailability (Schlieker et al., 2001; Vertačnik and Bišćan, 1993). Future research should aim to clarify the origin of REE concentration anomalies observed in certain lakes through targeted geochemical analyses. Additionally, the inclusion of both historically monitored TEs and emerging TCEs into long-term biomonitoring programs would improve risk assessments and provide useful information for lake management strategies.

#### 4.1.2. Trace element in fish liver

Our results demonstrate that anthropogenic inputs in the Rouyn-Noranda area have a widespread and systematic impact on aquatic ecosystems, affecting both abiotic (water and sediment) and biotic (fish) matrices. Consistent with patterns observed in water and sediment, TE concentrations in walleye and yellow perch liver were higher in lakes located close to the HS than in those further away. Pierron et al. (2009) also documented Cu concentrations in perch liver from N1 that were four times higher than those measured in nearby lake N2. Likewise, Giguère et al. (2004) reported significantly higher Cd concentrations in yellow perch from N1 and N2 compared to the more remote lake F1. Among the TEs, Cu, Se, and Cd were particularly assimilated in walleye, while Se was predominant in yellow perch, and their concentrations decreased significantly with increasing distance from the HS. In contrast, elements such as Zn, Pb, and As did not show significant spatial trends, despite being historically associated with mining activities in the region since the 1990s. These differences may reflect not only to species-specific homeostatic regulation mechanisms, but also to the complexity of environmental exposure pathways (Wang and Rainbow, 2008). For instance, if dietary uptake plays a significant role in TE accumulation, as suggested by Giguère et al. (2004), then direct relationships between water concentrations and tissue burdens may be obscured by trophic transfer. Moreover, because TE concentrations in water can fluctuate over time, the liver—being a long-term integrative organ—may reflect past rather than current exposure levels, further complicating spatial comparisons based solely on water chemistry (Giguère et al., 2005).

Interestingly, yellow perch generally exhibited higher concentrations of several TEs compared to walleye, including Cu, Pb, Zn, Se, As, Co, Tl, and Ti, across all six lakes. This pattern could be related to interspecific differences in feeding behavior, trophic position, and physiology. Yellow perch tend to feed closer to the benthic zone and may be more exposed to sediment-associated contaminants, whereas walleye are more pelagic and piscivorous, thereby occupying a higher trophic level (Blaney et al., in review). Trophic dilution can occur for certain TEs (e.g., Cu, As, Se), with lower concentrations observed in higher trophic-level fish (Chen et al., 2000; Griboff et al., 2018). Since walleye prey mainly on yellow perch in N1 and N3 (Blaney et al., in review), homeostatic regulation by metallothioneins may control metal uptake and excretion in biota, thereby limiting biomagnification (Nfon et al., 2009). In the case of As, this may also be explained by its low biomagnification potential (Asante et al., 2008).

As reported for abiotic matrices, some TEs (Co) and REEs (La, Ce) displayed opposite spatial patterns in livers of both fish species, with concentrations increasing with distance from the contamination source. These elements are typically considered to originate from natural sources, such as the erosion of REE-rich minerals in the local bedrock or surrounding soils (Ricard-Henderson, 2023). Furthermore, the La-Ce pair often co-occurs in natural environments due to their similar chemical properties and shared mineral hosts (Dushyantha et al., 2020). Their positive correlation with distance supports the hypothesis that they are not associated with smelting emissions but rather reflect background geochemical variability.

Together, these findings highlight the complex interplay between anthropogenic contamination and natural geochemical background in influencing the presence and distribution of trace elements in fish liver. While proximity to the contamination source remains the primary driver of enrichment for contaminants like Cu, Se, and Cd, other elements such as REEs and Co appear more influenced by natural variability associated to local geology. Understanding these dynamics is essential to accurately interpret bioaccumulation patterns and evaluate ecological risks.

### 4.2. Comparison of trace element levels in lake water and sediment with environmental guidelines

Elevated concentrations in sediments raise concerns about long-term accumulation and the potential remobilization of TE into the water column, particularly under climate change conditions. Climate change is increasing the frequency and intensity of processes that remobilize metals from sediments, especially through flooding and redox changes (Ciszewski and Grygar, 2016). Current environmental regulations often rely on measurements of total metal concentrations in water or sediments. However, this approach does not always reflect the actual ecological risk, since metal toxicity depends on local conditions, such as pH, organic matter, and oxygen levels, that influence the bioavailability fraction of metals to organisms (Väänänen et al., 2018).

Our results often exceeded the Canadian water and sediment quality guidelines for the protection of aquatic life, particularly for Cu, Zn, and Cd in water, and for Cu, Zn, Cd, As, and Pb in sediments. The concentrations of Cu, Zn, and Cd in water lake exceeded Canadian guidelines in N2 and N3, while in N1 only Cu exceeded the guideline. Overall, concentrations have declined compared to early 2000s measurements (Campbell et al., 2003) which may be explained by increased efforts to reduce industrial emissions and implement stricter environmental regulations since this period. Zn and Cd levels in N3 have dropped by 75% but still exceed water quality guidelines. For example, Cd in water is 2.5-fold the guideline. In contrast, Cu in N3 increased by over 65%, suggesting ongoing or new inputs from the HS, while both Cu and Cd decreased in N1, where Cu levels dropped nearly 2-fold (from 140 to 84 nmol/L), possibly due to hydrological changes following the separation of the southern and northern lake basins. Concentrations of As declined by about 15% in N1 over the past decade, remaining below water quality guidelines across all lakes (Proulx et al., 2015). Lakes I1 and F1 exhibited consistently low and stable concentrations of Cu, Zn, and Cd over time, indicating minimal metal inputs in these more remote ecosystems. Cd concentrations in F2 decreased by nearly 80% since 1989, now reaching levels approximately 75% below the CWQG, whereas Cu concentrations more than doubled over the same period, suggesting potential long-term contamination even in distant lakes (Perceval et al. 2006). This increase highlights that remote sites may also be vulnerable to delayed contamination effects. TEs such as Zn, Cu, Cd, Pb, and As often exceed national and international guidelines for the protection of aquatic life in water, especially in areas affected by mining, agriculture, or urbanization (Dengg et al., 2025; Edokpayi et al., 2016; Zhang et al., 2014). For instance, in a mining-impacted area of eastern China, Cu concentrations reached up to 133,700 nmol/L in river water near the source of contamination, gradually decreasing with distance from the mine (Zhang et al., 2014). Similarly, in the Mvudi River in South Africa, where municipal waste dumping has been identified as a major pollution source, Cu concentrations in water ranged from 378 to 2912 nmol/L (Edokpayi et al., 2016). In contrast, in undisturbed areas such as Lake Ōkataina in New Zealand, which is surrounded by native forest with no permanent human settlement in the watershed, Cu concentrations were considerably low, typically ranging from ∼0.1 to 0.4 nmol/L in surface and depth profiles, thereby providing a reference for background levels (Dengg et al., 2025). Our measured concentrations, ranging from 33 to 299 nmol/L across the six lakes, fall between these extremes.

Sediment TEs concentrations raise particular concern, as guideline exceedances were observed for several TEs across multiple lakes. When compared to historical data reported by Perceval et al. (2006b), our results suggest that guideline exceedances are not a new concern, but rather a persistent environmental issue in the region. For example, Cd and Cu concentrations in N1 and N3 declined compared to early 2000s values (e.g., Cd in N3 decreased by nearly 50%) but still exceed the CSQG (Pb in N1 surpasses current sediment guidelines by over 200%), indicating ongoing ecological risks. In I1 and F1, Cu, Zn, and Cd concentrations increased over time; for instance, in I1, Cd levels rose by 40% compared to 2000 values, exceeding the ISQG by nearly 480%, suggesting that even remote areas remain vulnerable to contamination. These parallel patterns in water and sediments underscore the chronic contamination in the Rouyn-Noranda region. Although improvements for some elements are evident, widespread exceedances persist, especially near historical sources, indicating that legacy pollution continues to affect these lake ecosystems. This situation reflects a broader national pattern, as a large-scale study of 167 Canadian lakes found that 70% of sites exceeded sediment quality guidelines for at least one toxic metal (Zilkey et al., 2025).

To the best of our knowledge, aqueous and sediment data for most TCEs and REEs are still very limited for the study lakes. Elements such as Ti, Sr, La, and Ce are increasingly detected in aquatic systems, yet no Canadian guidelines currently exist for them (Banaee et al., 2025; Hauser-Davis, 2023). Most existing studies focus on individual elements, and data on the occurrence and interactions of several elements and their ecological impacts (i.e., cocktail effect) are still scarce (Gu et al., 2023). These gaps highlight the need for targeted research and the development of specific regulatory benchmarks for these emerging contaminants. Even with some pioneer works regarding REE toxicity in aquatic environments, our current knowledge is not enough to better understand the impact of such TCEs.

### 4.3. Human risk assessment for fish consumption

While fish offer significant health benefits, food safety concerns, especially in regions with high fish consumption, require ongoing vigilance and regulation (Demelash Abera and Alefe Adimas, 2024). In our study, yellow perch and walleye are commonly consumed by local communities in the Abitibi region, making the evaluation of potential health risks from industrial contamination a priority. Muscle samples from both species exceeded fish consumption guidelines for at least one element (As, Zn, Cd or Pb) in all lakes. In several cases, multiple exceedances were observed within the same lake.

Interestingly, in Québec, public health advisories regarding fish consumption are primarily driven by mercury (Hg) concentrations, even though other contaminants (i.e., metals, polychlorinated biphenyls (PCBs), dioxins and furans, and polybrominated diphenyl ethers (PBDEs)) are also analyzed. However, these are generally present at low levels and are rarely the basis for consumption restrictions (Gouvernement du Québec, 2025). As a result, human populations that consume fish as a major part of their diet may be unknowingly exposed to dangerous levels of multiple contaminants. Moreover, recent evidence suggests that cooking can significantly alter TE concentrations in fish tissues, with changes varying by element and species. For example, As levels decreased by 14% while Se increased by 39% in whitefish after cooking, whereas other studies reported increases of up to 55% for Hg in marine mammals (Amyot et al., 2023). Such variability, driven by the chemical behavior and affinity of each element, challenges the validity of exposure models based solely on raw tissue concentrations. Muscle tissues of yellow perch and walleye from the Rouyn- Noranda area were last analyzed in 2009, when concentrations of As, Cd, Cu, Pb, and Zn were reported well below fish consumption guidelines (Proulx et al., 2015). In contrast, our results show a marked increase in TE concentrations over the past 15 years. For example, Cu concentrations in yellow perch from N1 increased from 0.3 mg/kg ww in 2009 to 7.7 mg/kg ww in our study. Similarly, Zn levels in walleye from N3 rose from 4.3 to 80 mg/kg ww, representing an increase of over 1760%. Increases in TE concentrations in fish muscle over time are mainly driven by persistent sediment contamination, remobilization of contaminants, ongoing human activities, physicochemical changes in the lake, and biological factors like species and age (Avigliano et al., 2015; Plessl et al., 2017). When compared with concentrations reported in yellow perch and walleye from other study areas, the levels observed in this study appear consistently higher. For instance, in Sudbury (Ontario)—a historically mining-impacted area—yellow perch showed Cu concentrations of 3.5 mg/kg ww, making the levels measured in perch from N1 approximately 2.2 times higher (Pyle et al., 2005). In that study, both Cu and Zn remained below consumption guidelines. In a low-impact area such as the Zlatar Reservoir in Serbia, pikeperch (*Sander lucioperca*) exhibited Cu concentrations around 0.97 mg/kg ww (Nikolić et al., 2022), nearly eight times lower than those we found in N1 perch.

Chronic exposure to TEs such as Cu, Zn, and Cd at levels exceeding WHO/FAO guidelines through fish consumption has been associated with adverse health effects in humans, including kidney dysfunction and an increased risk of cancer (Osman et al., 2025). Several studies found that estimated weekly intakes of Cd from fish can exceed safe limits, especially in populations with high fish consumption, posing a chronic health risk. Cu and Zn are essential nutrients, and most studies found that even when levels slightly exceed guidelines, the risk of adverse health effects is low for typical consumption rates. However, very high or prolonged exposure could cause toxicity, especially for Cu (Varol et al., 2020; Zafarzadeh et al., 2018).

We observed that high concentrations in fish were not always associated with the highest metal levels in water or sediment (as for As). This underscores the need to consider bioavailability, as total concentrations alone may not predict uptake in organisms (Väänänen et al., 2018). Furthermore, the lack of regulatory limits for many REEs and TCEs, such as La, Ce, Ti, and Tl, in fish muscle tissue represents a critical gap, especially given their increasing detection in freshwater ecosystems and unknown long-term health effects. Such data will support the development of more accurate guidelines that reflect both ecological and human health risks.

## 5. Conclusion

This study provides crucial insights into the complex and spatially variable distributions of TEs, including TCEs, in freshwater ecosystems impacted by historical mining and emerging e-waste recycling activities. It highlights distinct behaviors of historically reported versus emerging elements like Sr, Tl, Co, and REEs, emphasizing their different environmental sources and bioavailability throughout the aquatic environment and the food web. The detection of TCEs in aquatic biota—even in more distant lakes— raises important concerns about their bioavailability and ecological risks. By integrating chemical and biological data across multiple matrices, this research advances global understanding of contaminant dynamics in freshwater systems associated with mining-related operations. Given the growing industrial use and lack of regulation for many TCEs and REEs, these findings stress the urgent need to expand environmental monitoring and risk assessment frameworks worldwide to address emerging contaminants of increasing ecological and health significance. Future efforts should aim to establish threshold or guideline values for both abiotic (water, sediment) and biotic (organisms) matrices, investigate the ecological and geochemical mechanisms driving their distribution and bioaccumulation, and integrate TCEs and REEs into long-term environmental monitoring programs. Such multidisciplinary approaches are essential to better anticipate the environmental consequences of rising anthropogenic emissions and to support evidence-based management of freshwater ecosystems.

## CRediT authorship contribution statement

**Marta Gabriele:** Data Curation, Formal Analysis, Investigation, Methodology, Visualization, Writing – Original Draft, Writing – Review & Editing. **Guillaume Grosbois**: Supervision, Project Administration, Funding Acquisition, Conceptualization, Resources, Investigation, Methodology, Validation, Writing – Review & Editing. **Maikel Rosabal**: Supervision, Conceptualization, Resources, Investigation, Methodology, Validation, Writing – Review & Editing. **Miguel Montoro Girona:** Conceptualization, Funding Acquisition, Investigation, Resources, Supervision, Validation, Writing – Review & Editing. **Patrice Blaney:** Investigation, Writing – Review & Editing.

## Acknowledgments

We acknowledge the funds from the Hecla-mining company and the Interuniversity Group of Limnology Research (GRIL) for providing the necessary funding to support our project and the UQAT for supplying the essential equipment. We extend our gratitude to the interns and students from GREMA who participated in field collections and laboratory work: Ariane Barrette, Julianne Breton, Olivier Bruneau, Marilou Cournoyer, Justin Gagnon, Patrice Blaney, Éléa Jaskolski, Liv Jessen, Félix Labbé, Jade Lessard, Julie Marchal, Jérémy Mainville-Gamache, Chloé Tanguay, Antoine Villeneuve, and William Vincent. Additionally, we thank Olaloudé Judicaël Franck Ossé for assistance with statistical analyses. We also thank Technosub, OBVT, and the Collectif Territoire for their assistance throughout the project.

---

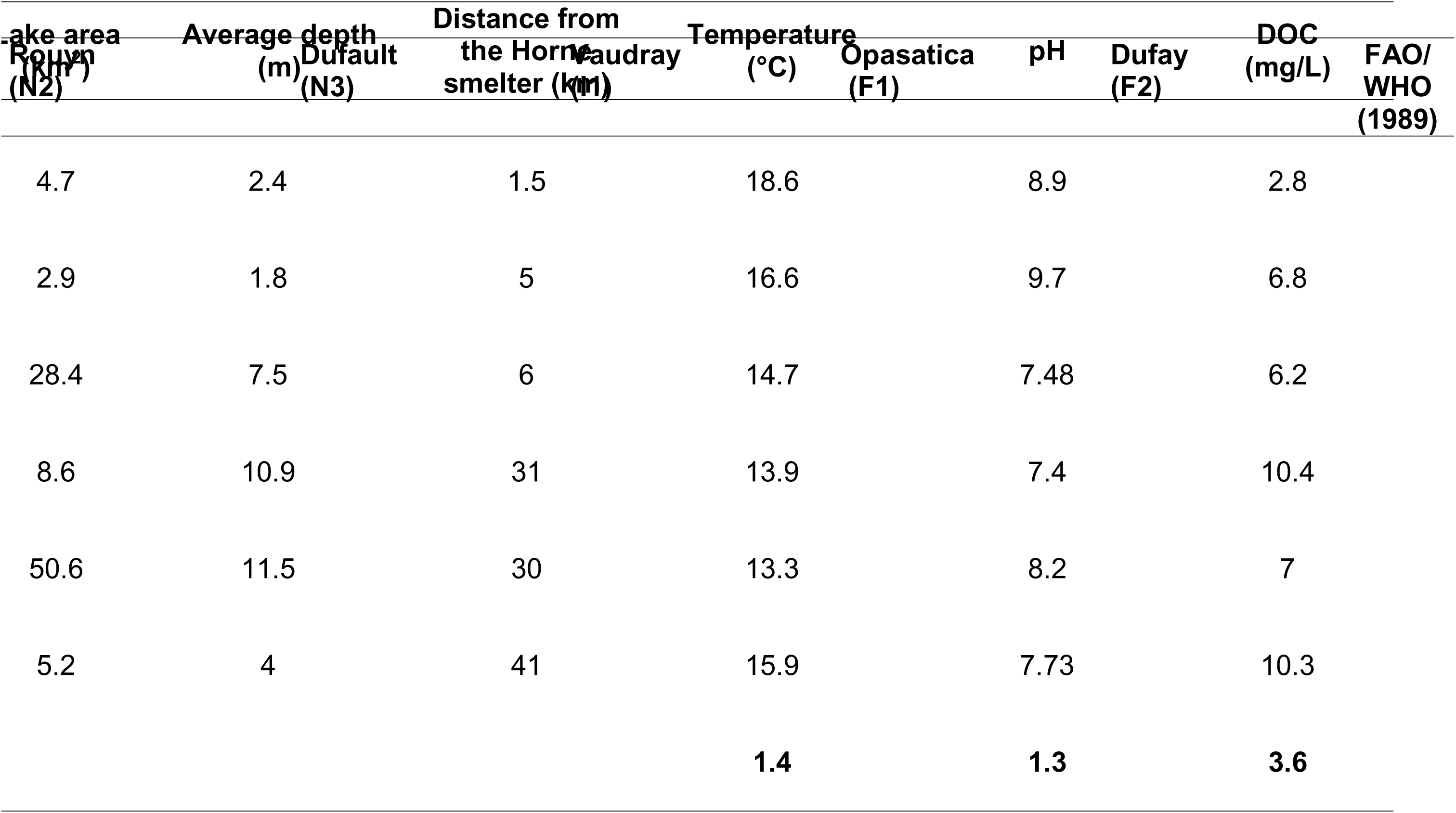
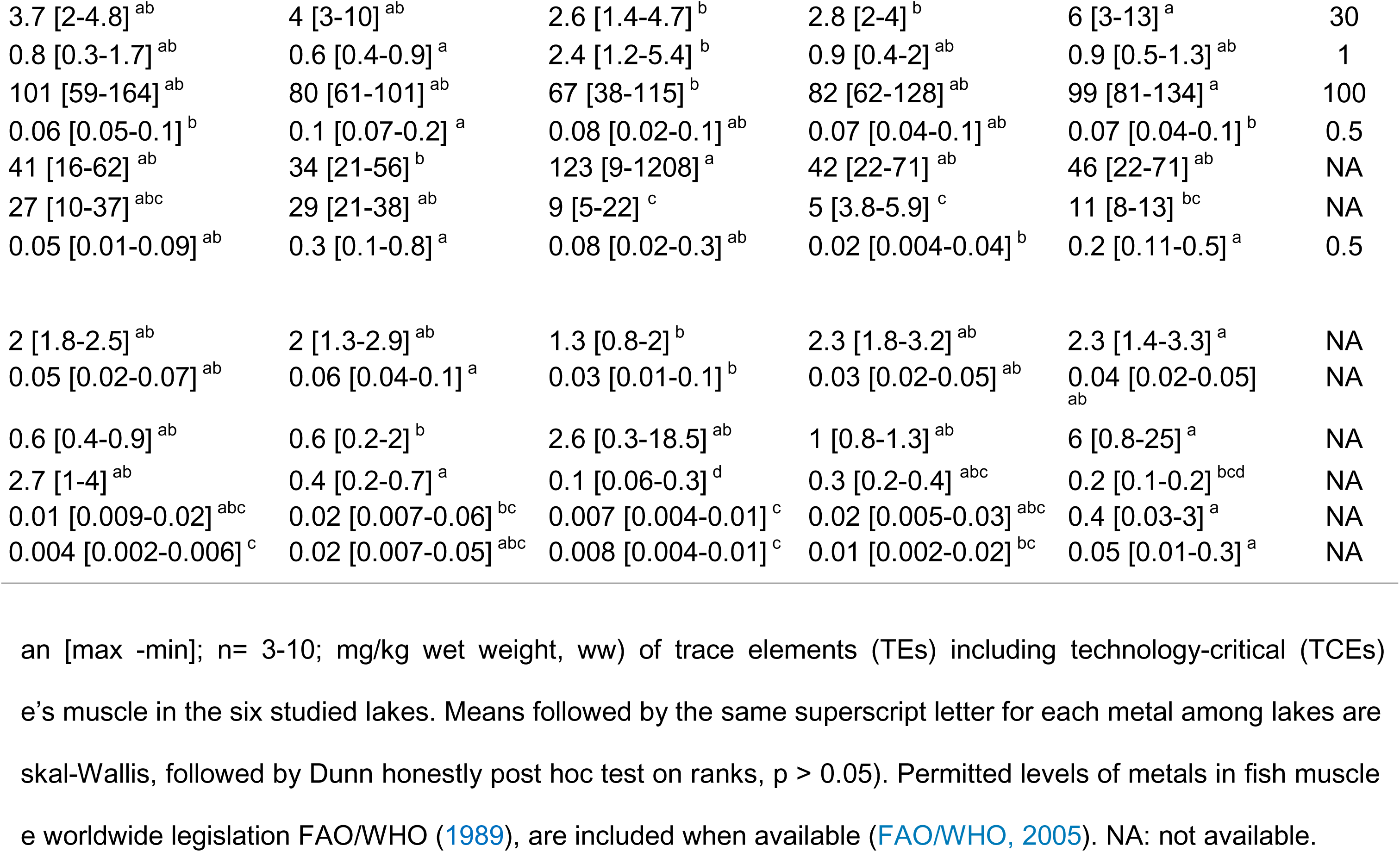
chemical characteristics of sampled lakes (DOC: dissolved organic carbon). Spatial gradient (calculated as the alue to the minimum mean value) for each characteristic is also given. Mean values are calculated from the three

|  | Rouyn<br>(N2) | Dufault<br>(N3) | Vaudray<br>(I1) | Opasatica<br>(F1) | Dufay<br>(F2) | FAO/WHO<br>(1989) |
| --- | --- | --- | --- | --- | --- | --- |
|  | 1.4 [1.3-1.5] | NA | 6.5 [3.4-13] | 1.3 [0.3-2.2] | 8.5 [4.7-13] | 30 |
|  | 0.06 [0.04-0.07]<sup>b</sup> | NA | 1 [0.5-1.8]<sup>a</sup> | 0.1 [0.03-0.2]<sup>b</sup> | 0.4 [0.3-0.5]<sup>ab</sup> | 1 |
|  | 176 [154-197] | NA | 144 [95-231] | 145 [130-159] | 133 [107-155] | 100 |
|  | 0.06 [0.05-0.07]<sup>b</sup> | NA | 0.5 [0.2-1.2]<sup>a</sup> | 0.05 [0.02-0.08]<sup>b</sup> | 1.3 [0.2-2.5]<sup>ab</sup> | 0.5 |
|  | 34 [32-36]<sup>b</sup> | NA | 67 [34-114]<sup>ab</sup> | 37 [14-60]<sup>ab</sup> | 162 [78-255]<sup>a</sup> | NA |
|  | 55 [54-56]<sup>ab</sup> | NA | 17 [7-23]<sup>bc</sup> | 6.3 [5.8-6.9]<sup>c</sup> | 22 [18-28]<sup>abc</sup> | NA |
|  | 0.4 [0.3-0.5] | NA | 1.7 [0.2-2.3] | 0.4 [0.3-0.5] | 3.7 [0.9-6.4] | 0.5 |
|  | 3.8 [3.3-4]<sup>a</sup> | NA | 2 [1.3-3.4]<sup>b</sup> | 5.3 [3.3-7.5]<sup>a</sup> | 3 [2-3.7]<sup>ab</sup> | NA |
|  | 0.3 [0.2-0.3]<sup>a</sup> | NA | 0.1 [0.05-0.3]<sup>ab</sup> | 0.2 [0.1-0.2]<sup>ab</sup> | 0.2 [0.1-0.3]<sup>ab</sup> | NA |
|  | 9 [4.8-13]<sup>ab</sup> | NA | 33 [1.3-114]<sup>a</sup> | 31 [24-38]<sup>a</sup> | 1.7 [1.4-2]<sup>ab</sup> | NA |
| }]<sup>b</sup> | 3.8 [3.5-4]<sup>a</sup> | NA | 0.1 [0.07-0.3]<sup>a</sup> | 0.2 [0.1-0.3]<sup>a</sup> | 0.1 [0.09-0.2]<sup>ab</sup> | NA |
|  | 0.02 [0.01-0.03]<sup>ab</sup> | NA | 0.02 [0.006-0.03]<sup>ab</sup> | 0.03 [0.02-0.05]<sup>ab</sup> | 0.07 [0.04-0.09]<sup>a</sup> | NA |
|  | 0.02 [0.007-0.03] | NA | 0.02 [0.003-0.04] | 0.03 [0.02-0.03] | 0.06 [0.05 - 0.07] | NA |

