## Supplementary Material for "Mining and e-waste recycling influence the spatial distribution of technology-critical elements, but not rare earth elements, in boreal lakes"

**SUPPLEMENTARY INFORMATION**

Table S1. Geographic location, distance (km), and orientation (east or west) from Horne smelter for each sampled lake (n=6). Sample types (water, sediments, and fish) collected is also indicated.

| **Lake**  **(CODE)** | **Location** | | **Distance from the Horne smelter (km)** | **Orientation (east or west)** | **Sample type** | | | | |
| --- | --- | --- | --- | --- | --- | --- | --- | --- | --- |
|  |  |  |  |  | **Water** | **Sediments** | | ***Perca flavences*** | ***Sander vitreus*** |
| Osisko (N1) | N48.2412; W79.01308 | | 1.5 | E | x | x | x | | x |
| Rouyn (N2) | N48.2389; W78.9542 | | 5 | E | x | x | x | | x |
| Dufault (N3) | N48.30709; W78.98737 | | 6 | E | x | x | x | | x |
| Vaudray (I1) | N48.09551; W78.67839 | | 31 | E | x | x | x | | x |
| Opasatica (F1) | | N48.1294; W79.3350 | 30 | W | x | x | x | | x |
| Dufay (F2) | N48.04098; W79.45187 | | 41 | W | x | x | x | | x |
|  | **Samples per site** | |  |  | 3 | 3-5 | 10 | | 10 |
|  | **Total samples** | |  |  | 18 | 20 | 41 | | 49 |

Table S2. Average, minimum and maximum physico-chemical characteristics of sampled lakes (DOC: dissolved organic carbon, DIC: dissolved inorganic carbon, POM: particulate organic matter, TP: total phosphorus, TN: total nitrogen). Spatial gradient calculated as the ratio of the maximum mean value to the minimum mean value for each characteristic is also given.

| **Lake (CODE)** | **Dissolved oxygen***  **(mg/L)** | | **Sp Cond***  **(µS/cm)** | | **DOC**^Ψ^  **(mg/L)** | **DIC**^Ψ^  **(mg/L)** | **POM^¥^**  **(mg/L)** | | **TP^**  **(µg/L)** | **TN^**  **(ppm)** |
| --- | --- | --- | --- | --- | --- | --- | --- | --- | --- | --- |
| **Osisko (N1)** | 8.5  [8. – 8.9] | 281  [278 - 361] | | 2.8  [2.7-2.8] | | 11.5  [10.8-12.1] | 1.8  [1.5-1.9] | | 20.1  [19.6-21] | 0.30  [0.22-0.35] |
| **Rouyn (N2)** | 8.7  [7.6 – 9.6] | 411  [383 - 657] | | 6.8  [6.8-6.8] | | 14.4  [14.1-14.9] | 9.3  [8.3-10.4] | | 130.6  [107.6-142.2] | 0.5  [0.4-0.6] |
| **Dufault (N3)** | 7.7  [4.6 – 10.9] | 140  [108 - 172] | | 6.2  [6-6.5] | | 5.7  [5.3-6.3] | 1.5  [1.5-1.6] | | 12.3  [9.9-16.7] | 0.2  [0.1-0.3] |
| **Vaudray (I1)** | 7.5  [3.3 – 8.5] | 27.7  [23.2 - 30.7] | | 10.4  [10-11] | | 1.9  [1.5-2.4] | 2.1  [1.9-2.3] | | 8  [7.6-8.5] | 0.2  [0.1-0.3] |
| **Opasatica (F1)** | 8.1  [6.4 – 10] | 77.2  [69.2 – 137] | | 7 | | 15.1 | 3.6  [3.5-3.6] | | 39.8  [17.8-67.2] | 0.3  [0.2-0.4] |
| **Dufay (F2)** | 7.6  [1.1 – 8.6] | 30.4  [26.8 - 54.2] | | 10.3  [9.9-10.6] | | 2.3  [2-2.5] | 2.8  [2.8-2.9] | | 22.3  [21.4-23.1] | 0.2  [0.1-0.3] |
| **Gradient** | **1.2** | **14.9** | | **3.6** | | **7.6** | **5.9** | **16.2** | | **2.8** |

* Water-column profiles of dissolved oxygen concentration (mg/L) and specific conductivity (µS/cm) were measured using a multiparameter profiler (RBR Concerto, Ottawa, ON). Water samples were retrieved from the epilimnion (0.5 m below the surface) at three distinct locations of the lake using a Ruttner water sampler and a bucket, previously cleaned with nitric acid (HNO_3_; omni-trace Grade) at 15% (v/v) and rinsed seven times with ultrapure water (Milli-Q system; >18 MΩ-cm) to avoid unintentional TE contamination. Ψ Water samples (n=3/lake) were filtered on 0.45 µm hydrophobic polytetrafluoroethylene membrane (Fisherbrand™, Pittsburgh, PA), gathered in brown pre-combusted vials (4h at 450 °C) and sent to the GRIL laboratory of the Université du Québec à Montréal (UQAM) for dissolved organic (DOC) analyses. DIC and DOC were analyzed with an OI Analytical Aurora 1030 W TOC Analyzer using a persulfate oxidation method. ¥ Additionally, 4 L of water was filtered through a pre-weighted Whatman glass fiber filter (0.7 μm pore size, 47 mm diameter, and 0.42 mm thickness) under moderate vacuum to collect POM. Filters were combusted at 60 °C for 24 h to remove the inorganic carbon and then at 450 °C for 4 h to remove OM. Finally, ^ water samples (n=3/lake) were collected in acid-washed (10% HCl) transparent 50 mL vials and sent to the GRIL laboratory of UQAM for total nitrogen (TN) and total phosphorus (TP) analyses. TP was quantified using the molybdenum-blue method following persulfate digestion at the GRIL-UQAM (Wetzel & Likens, 2000). TN was analyzed by an OI Analytic Flow Solution 3100, following a potassium persulfate digestion, coupled with a cadmium reactor.

Table S3. Morphological characteristics (mean ± standard deviation, SD) of adult yellow perch (*Perca flavescens*) and walleye (*Sander vitreus*) collected in six lakes of the Rouyn-Noranda mining region.

| **Lake**  **(CODE)** |  | **Yellow perch** | | | |  |  | **Walleye** | | | |
| --- | --- | --- | --- | --- | --- | --- | --- | --- | --- | --- | --- |
|  | Sex | N | Length (cm) | Weight  (g) | Condition factor |  | Sex | N | Length  (cm) | Weight  (g) | Condition factor |
| Osisko (N1) | F | 5 | 22 ± 0.96 | 125 ± 27 | 0.91 ± 0.53 |  | F | 5 | 41 ± 0.41 | 570 ± 42 | 0.79 ± 0.05 |
|  | M | 5 | 23 ± 1 | 113 ± 9 | 0.91 ± 0.06 |  | M | 5 | 42 ± 1 | 621± 51 | 0.78 ± 0.04 |
| Rouyn (N2) | NA | 2 | 10 ± 3 | 13 ± 11 | 0.99 ± 0.05 |  | NA | 3 | 51 ± 2 | 1455 ± 208 | 1 ± 0.06 |
| Dufault (N3) | NA | 9 | 11 ± 0.57 | 8.67 ± 1.23 | 0.64 ± 0.11 |  | F | 5 | 28 ± 6 | 185 ± 117 | 0.68 ± 0.10 |
|  |  |  |  |  |  |  | M | 5 | 33 ± 2 | 278 ± 49 | 0.76 ± 0.05 |
| Vaudray (I1) | F | 5 | 20 ± 3 | 95 ± 55 | 0.96 ± 0.19 |  | F | 5 | 32 ± 3 | 274 ± 73 | 0.76 ± 0.04 |
|  | M | 5 | 15 ± 2 | 44 ± 15 | 1 ± 0.16 |  | M | 5 | 30 ± 3 | 218 ± 72 | 0.78 ± 0.05 |
| Opasatica (F1) | NA | 2 | 11 ± 2 | 16 ± 9 | 1 ± 0.12 |  | NA | 6 | 35 ± 2 | 437 ± 100 | 1 ± 0.07 |
| Dufay (F2) | NA | 2 | 19 ± 3 | 76 ± 37 | 0.99 ± 0.035 |  | F | 5 | 29 ± 2 | 221 ± 49 | 0.82 ± 0.04 |
|  |  |  |  |  |  |  | M | 5 | 24 ± 1 | 119 ± 17 | 0.81 ± 0.02 |

Table S4. Recovery percentages (mean ± standard deviation; %) for trace elements (TEs, including technology-critical elements, TCEs) measured in BCR-668 (muscle tissue), DOLT-5 (dogfish liver), MESS-4 (marine sediment), TILL-3 (geochemical soil), BCR-667 (estuarine sediment), JSd-2 (stream sediment), and 1577b (bovine liver) as well as method detection limit (MDL) (µg/L) for trace elements using ICP-QQQ. Percentage of samples compared to the total samples numbers with concentrations higher than the MDL (%). Values with recovery percentages below 81% or over 126% are shown in bold.

|  |  |  |  |  |  |  |  |  | **% of samples with concentrations exceeding**  **the MDL relative to the total sample count** | | | |
| --- | --- | --- | --- | --- | --- | --- | --- | --- | --- | --- | --- | --- |
| **Trace element** | **BCR-668 (n=6-9)** | **DOLT-5**  **(n=3-5)** | **MESS-4**  **(n=6)** | **TILL-3**  **(n=6)** | **BCR-667**  **(n=6)** | **JSd-2**  **(n=6)** | **1577b**  **(n=3)** | **MDL** | **Water** |  | **Sediment** | **Fish** |
| **TE** |  |  |  |  |  |  |  |  |  |  |  |  |
| **Cu** |  | 103 ± 8 | 103 ± 8 | 102 ± 3 |  | 107 ± 2 | 120 ± 21 | 0.0008 | 100 |  | 100 | 100 |
| **As** |  | 87 ± 8 | 87 ± 2 | 93 ± 1 |  | 98 ± 2 |  | 0.0105 | 100 |  | 100 | 100 |
| **Zn** |  | 99 ± 8 | 107 ± 4 | 103 ± 5 |  |  | 110 ± 19 | 0.0935 | 100 |  | 100 | 100 |
| **Cd** |  | 93 ± 8 | 81 ± 10 | **70 ± 3** |  | 104 ± 3 | 114 ± 20  **(n = 3-5)** | 0.0018 | 100 |  | 100 | 100 |
| **Cr** |  |  | 81 ± 3 | **64 ± 1** |  | **61 ± 1** |  | 0.0185 | 100 |  | 100 | **64** |
| **Fe** |  | 95 ± 8 | 99 ± 3 |  |  |  | 109 ± 19 | 0.0021 | 100 |  | 100 | 100 |
| **Ni** |  |  | 97 ± 3 | 90 ± 1 |  | 97 ± 1 |  | 0.0075 | 100 |  | 100 | **56** |
| **Se** |  | 86 ± 10 |  |  |  | 126 ± 7 | 107 ± 22 | 0.0077 | 100 |  | 100 | 100 |
| **Mo** |  | **80 ± 7** |  |  |  | 101 ± 2 | 109 ± 17 | 0.0075 |  |  | 100 | **44** |
| **Ag** |  | 86 ± 10 | 124 ± 7 |  |  |  |  | 0.0036 |  |  | 90 | **44** |
| **Pb** |  | 103 ± 36 | 91 ± 3 | **67 ± 3** |  | 99 ± 2 |  | 0.0022 | 100 |  | 100 | 95 |
| **U** | 81 ± 10 | 83 ± 10 | **43 ± 3** | **59 ± 2** | **48 ± 1** |  |  | 0.0029 | 100 |  | 100 | **1** |
| **TCE** |  |  |  |  |  |  |  |  |  |  |  |  |
| **Ti** |  |  |  | **71 ± 6** |  |  |  | 0.0336 | 83 |  | 100 | 100 |
| **V** |  | 91 ± 7 | 84 ± 3 | **75 ± 1** |  |  |  | 0.0011 | 100 |  | 100 | **76** |
| **Co** |  | 89 ± 6 | 93 ± 2 | **74 ± 1** |  | 92 ± 1 |  | 0.0008 | 100 |  | 100 | 100 |
| **Sr** |  | 93 ± 8 | **69 ± 2** |  |  |  | 108 ± 20 | 0.0006 | 100 |  | 100 | 100 |
| **Y** | 88 ± 12 |  |  | **59 ± 1** |  |  |  | 0.0012 | 100 |  | 100 | **71** |
| **Ru** |  |  |  |  |  |  |  | 0.0399 | **0** |  | **0** | **0** |
| **Pd** |  |  |  |  |  | **0** |  | 0.0084 | **0** |  | 100 | **0** |
| **Sb** |  | 90 ± 21 |  | **70 ± 3** |  |  |  | 0.0043 | 100 |  | 100 | **18** |
| **Pt** |  |  |  |  |  | **23392± 10^9^** |  | 0.0034 | **6** |  | **0** | **0** |
| **W** |  |  |  |  |  |  |  | 0.7491 | **0** |  | **0** | **0** |
| **Tl** |  |  | **73 ± 3** |  |  |  |  | 0.0009 | 100 |  | 100 | 89 |
| **La** | 88 ± 11 |  |  | 83 ± 3 | **33 ± 1** | 87 ± 3 |  | 0.0004 | 100 |  | 100 | 100 |
| **Ce** | 87 ± 14 |  |  | 86 ± 3 | **135 ± 2** | 83 ± 3 |  | 0.0007 | 100 |  | 100 | 89 |
| **Pr** | 85 ± 14 |  |  |  | **73 ± 1** | 106 ± 3 |  | 0.0004 | 100 |  | 100 | **51** |
| **Nd** | 94 ± 16 |  |  | 106 ± 4 | **74 ± 1** | 84 ± 2 |  | 0.0016 | 100 |  | 100 | **33** |
| **Sm** | 81 ± 11 |  |  | 95 ± 4 | 89 ± 2 | 96 ± 3 |  | 0.0015 | 88 |  | 100 | **8** |
| **Eu** | 92 ± 21 |  |  |  | 86 ± 2 | 92 ± 3 |  | 0.0012 | 88 |  | 100 | **0** |
| **Gd** | 102 ± 4 |  |  |  | 93 ± 1 | 99 ± 2 |  | 0.0011 | 88 |  | 100 | **26** |
| **Tb** | **76 ± 26** |  |  |  | **79 ± 2** |  |  | 0.0007 | 88 |  | 100 | **0** |
| **Dy** | 84 ± 14 |  |  |  | **76 ± 1** | 86 ± 2 |  | 0.0008 | 88 |  | 100 | **8** |
| **Ho** |  |  |  |  | **66 ± 1** | **69 ± 1** |  | 0.0003 | 88 |  | 100 | **4** |
| **Er** | 92 ± 11 |  |  | **73 ± 2** | **63 ± 2** | 93 ± 4 |  | 0.0012 | 88 |  | 100 | **0** |
| **Tm** | 102 ± 4 |  |  |  | **55 ± 2** |  |  | 0.0004 | 88 |  | 100 | **1** |
| **Yb** |  |  |  | **61 ± 3** | **53 ± 1** | **0** |  | 0.0011 | 88 |  | 100 | **1** |
| **Lu** |  |  |  | **59 ± 3** | **44 ± 2** | **51 ± 1** |  | 0.0162 | 88 |  | 100 | **0** |

Table S5. Spatial gradient ([TE]_maximum_/[TE]_minimum_) for each TE estimated from abiotic (water and sediments) and biotic (yellow perch and walleye muscle and liver) samples.

| **Trace**  **elements** | **Gradient** | | | | | | | |
| --- | --- | --- | --- | --- | --- | --- | --- | --- |
|  | **Water** | **Sediments** | **Yellow perch**  **(liver)** | | **Walleye (liver)** | **Yellow perch**  **(muscle)** | | **Walleye**  **(muscle)** |
| **TE** |  |  |  |  | |  | | |
| **Cu** | 9 | 160 | 15 | 4 | | 271 | 3 | |
| **As** | 5 | 78 | 7 | 5 | | 17 | 6 | |
| **Se** | 10 | 183 | 4 | 6 | | 9 | 8 | |
| **Cd** | 31 | 110 | 18 | 5 | | 39 | 3 | |
| **Cr** | 22 | 2 |  |  | |  |  | |
| **Fe** | 10 | 5 | 11 | 5 | | 716 | 4 | |
| **Ni** | 6 | 2 |  |  | |  |  | |
| **Zn** | 38 | 42 | 2 | 2 | | 8 | 2 | |
| **Pb** | 12 | 7 | 18 | 4 | | 10 | 29 | |
| **U** | 8 | 3 |  |  | |  |  | |
| **Mo** |  | 59 |  |  | |  | | |
| **Ag** |  | 268 |  |  | |  | | |
| **TCE** |  |  |  |  | |  | | |
| **Ti** | 138 | 2 | 5 | 9 | | 3 | 2 | |
| **V** | 35 | 1 |  |  | |  | | |
| **Co** | 23 | 5 | 25 | 3 | | 523 | 2 | |
| **Sr** | 6 | 1 | 34 | 9 | | 19 | 10 | |
| **Y** | 149 | 2 |  |  | |  | | |
| **Pd** |  | 3 |  |  | |  | | |
| **Sb** | 33 | 213 |  |  | |  | | |
| **Pt** | 17 |  |  |  | |  | | |
| **W** |  | 40 |  |  | |  | | |
| **Tl** | 138 | 168 | 161 | 18 | | 86 | 22 | |
| **La** | 328 | 2 | 74 | 24 | | 11 | 50 | |
| **Ce** | 386 | 1 | 88 | 17 | | 12 | 12 | |
| **Pr** | 233 | 1 |  |  | |  | | |
| **Nd** | 295 | 1 |  |  | |  | | |
| **Sm** | 215 | 2 |  |  | |  | | |
| **Eu** | 14 | 1 |  |  | |  | | |
| **Gd** | 67 | 2 |  |  | |  | | |
| **Tb** | 39 | 2 |  |  | |  | | |
| **Dy** | 76 | 2 |  |  | |  | | |
| **Ho** | 49 | 2 |  |  | |  | | |
| **Er** | 79 | 2 |  |  | |  | | |
| **Tm** | 18 | 2 |  |  | |  | | |
| **Yb** | 94 | 2 |  |  | |  | | |
| **Lu** | 24 | 1 |  |  | |  | | |

Table S6. Concentration (mean ± standard deviation, SD) of trace elements (TEs) measured in water (n= 3; nmol/L) in the six studied lakes. Water across the lake was collected in different locations. The spatial gradient ([TE]_maximum_/[TE]_minimum_) for each TE estimated from water and the Canadian Water Quality Guidelines (CWQGs) (nmol/L) are included when available (CCME, 2007). Letters are different when significant differences exist between means (Kruskal-Wallis, followed by Dunn honestly post hoc test on ranks, *p* < 0.05).

| **Lake**  **(CODE)** | **Osisko**  **(N1)** | **Rouyn**  **(N2)** | **Dufault**  **(N3)** | **Vaudray**  **(I1)** | **Opasatica**  **(F1)** | **Dufay**  **(F2)** | **CWQGs** | **Gradient** |
| --- | --- | --- | --- | --- | --- | --- | --- | --- |
| **Cu** | 89 ± 2 ^ab^ | 190 ± 0.8 ^ab^ | 299 ± 5 ^a^ | 47 ± 0.58 ^ab^ | 35 ± 0.1 ^ab^ | 33 ± 0.1 ^b^ | **47** | **9** |
| **As** | 23 ± 0.44 ^ab^ | 30 ± 0.55 ^a^ | 16 ± 0.29 ^ab^ | 13 ± 0.21 ^ab^ | 6.5 ± 0.09 ^b^ | 8.5 ± 0.08 ^ab^ | **67** | **5** |
| **Se** | 8 ± 1 ^ab^ | 13 ± 0.5 ^a^ | 7 ± 0.17 ^ab^ | 2 ± 0.12 ^ab^ | 1.3 ± 0.24 ^b^ | 1.6 ± 0.06 ^ab^ | **-** | **10** |
| **Cd** | 0.31 ± 0.03 ^ab^ | 1.3 ± 0.02 ^ab^ | 2 ± 0.06 ^a^ | 0.6 ± 0.01 ^ab^ | 0.07 ± 0.003 ^b^ | 0.2 ± 0.007 ^ab^ | **0.8** | **31** |
| **Cr** | 0.59 ± 0.19 ^a^ | 6 ± 0.09 ^ab^ | 2.4 ± 0.22 ^ab^ | 12.40 ± 0.10 ^ab^ | 4.73 ± 0.16 ^ab^ | 12 ± 0.09 ^b^ | **19** | **22** |
| **Fe** | 417 ± 107 ^a^ | 3909 ± 102 ^ab^ | 1662 ± 73 ^ab^ | 3261 ± 17 ^ab^ | 1775 ± 13 ^ab^ | 4476 ± 37 ^b^ | **-** | **10** |
| **Ni** | 12 ± 0.3 ^ab^ | 57 ± 0.20 ^a^ | 9 ± 0.22 ^b^ | 15 ± 0.22 ^ab^ | 10 ± 0.09 ^ab^ | 13 ± 0.08 ^ab^ | **-** | **6** |
| **Zn** | 86 ± 15 ^ab^ | 317 ± 9 ^ab^ | 528 ± 4 ^a^ | 72 ± 12 ^ab^ | 14 ± 3 ^b^ | 31 ± 8 ^ab^ | **107** | **38** |
| **Pb** | 0.2 ± 0.01 ^a^ | 2.6 ± 0.1 ^b^ | 1.7 ± 0.06 ^ab^ | 1 ± 0.2 ^ab^ | 0.27 ± 0.01 ^ab^ | 1.2 ± 0.03 ^ab^ | **-** | **12** |
| **U** | 0.06 ± 0.003 ^ab^ | 0.34 ± 0.006 ^a^ | 0.08 ± 0.002 ^ab^ | 0.05 ± 0.003^b^ | 0.22 ± 0.004 ^ab^ | 0.22 ± 0.003 ^ab^ | **63** | **8** |

Table S7. Concentration (mean ± standard deviation) of technology-critical elements measured in water (n= 3; nmol/L) in the six studied lakes. Water was collected in different locations across the lake. The spatial gradient ([TE]_maximum_/[TE]_minimum_) for each REEs estimated from water and the background concentrations in surface water (nmol/L) for REEs proposed by the National Institute of Health and the Environment in the Netherlands (nmol/L) are included. * Background concentrations were obtained from (Sneller et al., 2000) and represent detection limits reported for relatively pristine surface waters, as measured concentrations were below these limits (Van Son, 1994). Letters that are different indicate significant differences between means (Kruskal-Wallis, followed by Dunn honestly post hoc test on ranks, *p* < 0.05).

| **Lake**  **(CODE)** | **Osisko**  **(N1)** | **Rouyn**  **(N2)** | **Dufault**  **(N2)** | **Vaudray**  **(I1)** | | | **Opasatica**  **(F1)** | **Dufay**  **(F2)** | **Background concentrations*** | **Gradient** |
| --- | --- | --- | --- | --- | --- | --- | --- | --- | --- | --- |
| **Ti** | 0.6 ± 0.08 ^a^ | 37 ± 0.3 ^ab^ | 9.8 ± 0.5 ^ab^ | | 31 ± 0.2 ^ab^ | | 28 ± 0.9 ^ab^ | 82 ± 1 ^b^ | **-** | **138** |
| **Co** | 0.6 ± 0.04 ^a^ | 14 ± 0.11 ^b^ | 1.8 ± 0.03 ^ab^ | | 1.3 ± 0.03 ^ab^ | | 2.7 ± 0.003 ^ab^ | 2.3 ± 0.05 ^ab^ | **-** | **23** |
| **Sr** | 794 ± 17 ^ab^ | 904 ± 5 ^a^ | 290 ± 5 ^ab^ | | 147 ± 0.9 ^ab^ | | 419 ± 2 ^ab^ | 144 ± 1 ^b^ | **-** | **6** |
| **Tl** | 0.04 ± 0.003 ^ab^ | 2 ± 0.01 ^a^ | 0.04 ± 0.001 ^ab^ | | 0.02 ± 0.001 ^ab^ | | 0.01 ± 0.0002 ^b^ | 0.03 ± 0.001 ^ab^ | **-** | **138** |
| **La** | 0.01 ± 0.0003 ^a^ | 1.8 ± 0.02 ^ab^ | 0.4 ± 0.01 ^ab^ | | 1.3 ± 0.01 ^ab^ | | 0.97 ± 0.01 ^ab^ | 3.6 ± 0.04 ^b^ | **<0.57** | **328** |
| **Ce** | 0.02 ± 0.001 ^a^ | 3 ± 0.05 ^ab^ | 0.7 ± 0.01 ^ab^ | | 2 ± 0.02 ^ab^ | | 1 ± 0.02 ^ab^ | 6 ± 0.05 ^b^ | **<0.92** | **386** |
| **V** | 0.36 ± 0.02 ^a^ | 8 ± 0.07 ^ab^ | 2 ± 0.02 ^ab^ | | 4 ± 0.05 ^ab^ | | 6 ± 0.03 ^ab^ | 12 ± 0.24 ^b^ | **-** | **35** |
| **Y** | 0.02 ± 0.001 | 1.56 ± 0.02 | 0.4 ± 0.01 | | 1.2 ± 0.01 | | 0.98 ± 0.01 | 2.8 ± 0.03 | **<2.47** | **148** |
| **Sb** | 4 ± 0.10 ^ab^ | 13 ± 0.07 ^a^ | 2 ± 0.09 ^ab^ | | 0.53 ± 0.02 ^ab^ | | 0.4 ± 0.01 ^b^ | 0.39 ± 0.006 ^b^ | **-** | **33** |
| **Pt** | 0.003 ± 0.001 ^ab^ | 0.002 ± 0.0002 ^ab^ | 0.002 ± 0.0002 ^ab^ | | 0.001 ± 0.0007^ab^ | | 0.002 ± 0.001 ^ab^ | 0.0007 ± 0.001^b^ | **-** | **17** |
| **Pr** | 0.004 ± 0.0004 ^a^ | 0.46 ± 0.01 ^ab^ | 0.11 ± 0.004 ^ab^ | | 0.35 ± 0.001 ^ab^ | | 0.24 ± 0.003 ^ab^ | 0.88 ± 0.01 ^b^ | **<0.57** | **233** |
| **Nd** | 0.011 ± 0.001 ^a^ | 1.8451 ± 0.01 ^ab^ | 0.44 ± 0.01 ^ab^ | | 1.4 ± 0.002 ^ab^ | | 1 ± 0.02 ^ab^ | 3 ± 0.004 ^b^ | **<3** | **295** |
| **Sm** | 0.003 ± 0.002 ^a^ | 0.3003 ± 0.02 ^ab^ | 0.07 ± 0.01 ^ab^ | | 0.23 ± 0.01 ^ab^ | | 0.2 ± 0.004 ^ab^ | 0.5 ± 0.01 ^b^ | **<4** | **215** |
| **Gd** | 0.01 ± 0.0013 ^a^ | 0.3 ± 0.03 ^ab^ | 0.06 ± 0.01 ^ab^ | | 0.19 ± 0.003 ^ab^ | | 0.14 ± 0.004 ^ab^ | 0.4 ± 0.01 ^b^ | **<2.09** | **67** |
| **Dy** | 0.004 ± 0.0003 ^a^ | 0.16 ± 0.003 ^ab^ | 0.05 ± 0.004 ^ab^ | | 0.13 ± 0.01 ^ab^ | | 0.1 ± 0.01 ^ab^ | 0.3 ± 0.001 ^b^ | **<1.35** | **76** |
| **Yb** | 0.002 ± 0.001 ^a^ | 0.08 ± 0.0095 ^ab^ | 0.02 ± 0.004 ^ab^ | | 0.05 ± 0.003 ^ab^ | | 0.0504 ± 0.005 ^ab^ | 0.1 ± 0.01 ^b^ | **<0.75** | **94** |
| **Eu** | 0.01 ± 0.002 ^a^ | 0.0579 ± 0.004 ^ab^ | 0.02 ± 0.002 ^ab^ | | 0.05 ± 0.01 ^ab^ | | 0.03 ± 0.002 ^ab^ | 0.09 ± 0.004 ^b^ | **<1** | **14** |
| **Tb** | 0.001 ± 0.0001 ^a^ | 0.0305 ± 0.001 ^ab^ | 0.01 ± 0.001 ^ab^ | | 0.05 ± 0.003 ^ab^ | | 0.02 ± 0.001 ^ab^ | 0.05 ± 0.002 ^b^ | **<0.63** | **39** |
| **Ho** | 0.001 ± 0.0001 ^a^ | 0.03 ±0.0003 ^ab^ | 0.01 ±0.001 ^ab^ | | 0.02 ±0.001 ^ab^ | | 0.0233 ± 0.001 ^ab^ | 0.06 ± 0.001 ^b^ | **<0.54** | **49** |
| **Er** | 0.002 ± 0.0003 ^a^ | 0.08± 0.003 ^ab^ | 0.03 ± 0.004 ^ab^ | | | 0.07 ± 0.001 ^ab^ | 0.06 ± 0.005 ^ab^ | 0.16 ± 0.004 ^b^ | **<0.82** | **79** |
| **Tm** | 0.001± 0.0001 ^a^ | 0.01± 0.0007 ^ab^ | 0.005 ± 0.0001 ^ab^ | | | 0.01 ± 0.0005 ^ab^ | 0.009 ± 0.0007 ^ab^ | 0.02± 0.001 ^b^ | **<0.40** | **18** |
| **Lu** | 0.001 ± 0.0001 ^a^ | 0.013 ± 0.001 ^ab^ | 0.004 ± 0.001 ^ab^ | | | 0.01 ± 0.001 ^ab^ | 0.008 ± 0.001 ^ab^ | 0.02 ± 0.001 ^b^ | **<0.28** | **24** |

Table S8. Concentration (mean ± standard deviation) of trace elements measured in sediments (n= 3-5; µg/g dry weight, dw; 0-5 cm) in the six studied lakes. Sediments across the lake were collected in different locations. The spatial gradient ([TE]_maximum_/[TE]_minimum_) for each TE estimated from sediment and the Canadian sediment quality guidelines (CSQGs; µg/g dw) are included when available. Letters that are different indicate significant differences between means (Kruskal-Wallis, followed by Dunn honestly post hoc test on ranks, *p* < 0.05).

| **Lake**  **(Code)** | **Osisko**  **(N1)** | **Rouyn**  **(N2)** | **Dufault**  **(N3)** | **Vaudray**  **(I1)** | **Opasatica**  **(F1)** | **Dufay**  **(F2)** | **CSQGs** | **Gradient** |
| --- | --- | --- | --- | --- | --- | --- | --- | --- |
| **Cu** | 4504 ± 599 ^a^ | 1371 ± 504 ^ab^ | 1184 ± 384 ^ab^ | 49 ± 43 ^ab^ | 45 ± 20 ^ab^ | 28 ± 2 ^b^ | **36** | **160** |
| **As** | 227 ± 42 ^a^ | 73 ± 16 ^ab^ | 123 ± 30 ^ab^ | 35 ± 40 ^ab^ | 2.9 ± 1.2 ^b^ | 5 ± 1.5 ^ab^ | **6** | **78** |
| **Se** | 75 ± 22 ^a^ | 35 ± 9 ^ab^ | 22 ± 9 ^ab^ | 1.44 ± 1 ^ab^ | 0.41 ± 0.25 ^b^ | 0.56 ± 0.03 ^b^ | **-** | **183** |
| **Cd** | 77 ± 20 ^a^ | 55 ± 17 ^ab^ | 29 ± 8 ^ab^ | 3.5 ± 3 ^ab^ | 0.7 ± 0.67 ^b^ | 1.25 ± 0.28 ^ab^ | **0.6** | **110** |
| **Cr** | 62 ± 7 | 74 ± 7 | 62 ± 0.5 | 34 ± 22 | 91 ± 10 | 82 ± 1 | **37** | **2** |
| **Fe** | 126 10^3^ ± 30 10^3 a^ | 45 10^3^ ± 11 10^3 ab^ | 51 10^3^ ± 5 10^3 ab^ | 60 10^3^ ± 75 10^3 ab^ | 27 10^3^ ± 2 10^3 ab^ | 23 10^3^ ± 2 10^3 b^ | **-** | **5** |
| **Ni** | 84 ± 6 | 68 ± 5 | 43 ± 1 | 35 ± 64 | 47 ± 4 | 48 ± 2 | **-** | **2** |
| **Zn** | 4705 ± 1587 ^a^ | 1576 ± 314 ^ab^ | 2088 ± 365 ^ab^ | 214 ± 168 ^ab^ | 111 ± 39 ^b^ | 140 ± 11 ^b^ | **123** | **42** |
| **Pb** | 107 ± 14 ^ab^ | 143 ± 9 ^a^ | 72 ± 16 ^ab^ | 62 ± 72 ^ab^ | 20 ± 13 ^b^ | 33 ± 7.5 ^ab^ | **35** | **7** |
| **U** | 1.16 ± 0.17 ^ab^ | 0.96 ± 0.27 ^ab^ | 1.23 ± 0.07 ^ab^ | 0.48 ± 0.32 ^b^ | 1.66 ± 0.22 ^a^ | 1.18 ± 0.14 ^ab^ | **-** | **3** |
| **Ag** | 20 ± 2 ^a^ | 3.36 ± 5 ^ab^ | 5 ± 1.4 ^ab^ | 0.16 ± 0.22 ^b^ | 0.07 ± 0.07 ^b^ | 0.13 ± 0.02 ^ab^ | **-** | **268** |
| **Mo** | 12 ± 3 | 19 ± 22 | 2 ± 0.25 | 1 ± 5 | 0.42 ± 0.46 | 0.45 ± 0.05 | **-** | **59** |

| **Lake**  **(Code)** | **Osisko**  **(N1)** | **Rouyn**  **(N2)** | **Dufault**  **(N3)** | **Vaudray**  **(I1)** | **Opasatica**  **(F1)** | **Dufay**  **(F2)** | **Background concentrations** | **Gradient** |
| --- | --- | --- | --- | --- | --- | --- | --- | --- |
| **Ti** | 1087 ± 49 ^a^ | 1608 ± 88 ^ab^ | 1349 ± 63 ^ab^ | 854 ± 293 ^a^ | 1918 ± 62 ^b^ | 1710 ± 64 ^ab^ | **-** | **2** |
| **Co** | 85 ± 38 | 30 ± 3 | 79 ± 16 | 32 ± 38 | 16 ± 0.54 | 17 ± 1 | **-** | **5** |
| **Sr** | 38 ± 13 | 52 ± 3 | 39 ± 4 | 36 ± 8 | 52 ± 2 | 46 ± 5 | **-** | **1** |
| **Tl** | 1 ± 0.2 ^ab^ | 34 ± 9 ^a^ | 0.6 ± 0.1 ^ab^ | 0.2 ± 0.1 ^b^ | 0.3 ± 0.05 ^ab^ | 0.5 ± 0.1 ^ab^ | **-** | **168** |
| **La** | 25 ± 3 | 25 ± 1 | 28 ± 1 | 18 ± 16 | 31 ± 1.5 | 26 ± 2 | **37** | **2** |
| **Ce** | 50 ± 6 | 50 ± 2 | 58 ± 4 | 45 ± 41 | 62 ± 2.8 | 55 ± 5 | **69** | **1** |
| **V** | 43 ± 6 | 57 ± 5 | 58 ± 0.3 | 58 ± 26 | 61 ± 2 | 60 ± 0.51 | **-** | **1** |
| **Y** | 17 ± 3 | 13 ± 0.48 | 14 ± 1.3 | 9 ± 5 | 12 ± 0.25 | 13 ± 1 | **-** | **2** |
| **Sb** | 16 ± 1 ^a^ | 20 ± 5 ^a^ | 3 ± 2 ^ab^ | 0.47 ± 0.28 ^ab^ | 0.06 ± 0.05 ^b^ | 0.14 ± 0.05 ^b^ | **-** | **213** |
| **W** | 6 ± 2 ^a^ | 1.7 ± 1.4 ^ab^ | 0.33 ± 0.1 ^ab^ | 0.06 ± 0.4 ^b^ | 0.07 ± 0.06 ^b^ | 0.17 ± 0.05 ^a^ | **-** | **40** |
| **Pr** | 6 ± 0.89 | 5.9 ± 0.18 | 6.3 ± 0.41 | 5 ± 4 | 7 ± 0.24 | 6 ± 0.46 | **8** | **1** |
| **Nd** | 24.5 ± 3.7 | 22 ± 0.58 | 24 ± 1.6 | 18 ± 14 | 26 ± 0.68 | 23 ± 1.78 | **36** | **1** |
| **Sm** | 5 ± 0.8 | 4.22 ± 0.13 | 4 ± 0.35 | 3.23 ± 2 | 4.5 ± 0.12 | 4.12 ± 0.31 | **6** | **2** |
| **Gd** | 4.7 ± 0.8 | 3.64 ± 0.12 | 3.6 ± 0.32 | 2.73 ± 2 | 3.6 ± 0.10 | 3.4 ± 0.3 | **5** | **2** |
| **Dy** | 3.4 ± 0.5 | 2.6 ± 0.10 | 2.7 ± 0.3 | 1.93 ± 1.37 | 2.5 ± 0.02 | 2.4 ± 0.2 | **4** | **2** |
| **Yb** | 1.3 ± 0.16 | 1.1 ± 0.06 | 1.1 ± 0.10 | 0.80 ± 0.52 | 1.14 ± 0.03 | 1.07 ± 0.1 | **1.6** | **2** |
| **Eu** | 0.9 ± 0.15 | 0.8 ± 0.03 | 0.8 ± 0.08 | 0.69 ± 0.49 | 0.84 ± 0.03 | 0.79 ± 0.07 | **1** | **1** |
| **Tb** | 0.6 ± 0.09 | 0.4 ± 0.02 | 0.4 ± 0.06 | 0.34 ± 0.25 | 0.47 ± 0.02 | 0.44 ± 0.04 | **0.70** | **2** |
| **Ho** | 0.6 ± 0.08 | 0.4 ± 0.03 | 0.5 ± 0.06 | 0.35 ± 0.23 | 0.49 ± 0.01 | 0.46 ± 0.04 | **0.68** | **2** |
| **Er** | 1.70 ± 0.22 | 1.3 ± 0.08 | 1.3 ± 0.15 | 0.96 ± 0.62 | 1.36 ± 0.06 | 1.2 ± 0.17 | **4** | **2** |
| **Tm** | 0.22 ± 0.02 | 0.18 ± 0.01 | 0.2 ± 0.02 | 0.14 ± 0.08 | 0.19 ± 0.003 | 0.18 ± 0.02 | **0.26** | **2** |
| **Lu** | 0.19 ± 0.18 | 0.17 ± 0.15 | 0.17 ± 0.12 | 0.18 ± 0.03 | 0.20 ± 0.03 | 0.24 ± 0.03 | **0.22** | **1** |
| **Pd** | 0.16 ± 0.02 ^a^ | 0.08 ± 0.02 ^ab^ | 0.06 ± 0.01 ^b^ | 0.06 ± 0.03 ^ab^ | 0.07 ± 0.01 ^ab^ | 0.07 ± 0.02 ^ab^ | **-** | **3** |

Table S9. Concentration (mean ± standard deviation, sd) of technology-critical elements measured in sediments (n= 3-5; µg/g dry weight, w; 0-5 cm) in the six studied lakes. The spatial gradient ([TE]_maximum_/[TE]_minimum_) for each REEs estimated from sediment and the background concentrations in sediment for REEs proposed by the National Institute of Health and the Environment in the Netherlands (µg/g dry weight, dw) are included. * Background concentrations were obtained from (Sneller et al., 2000) and represent detection limits reported for relatively pristine surface waters, as measured concentrations were below these limits (Van Son, 1994). Letters that are different indicate significant differences between means (Kruskal-Wallis, followed by Dunn honestly post hoc test on ranks, *p* < 0.05).

Table S10. Summary of linear regression analyses of trace element (TE) concentrations in lake water and sediments as a function of distance from the Horne smelter. Regression coefficients (β), R² values, and significance levels are reported. Data from all sampled lakes were included for each element (n=3/lake).

| **Water** |  |  |  |  | **Sediment** |  |  |  |
| --- | --- | --- | --- | --- | --- | --- | --- | --- |
| **Element** | **Coefficient β** | **R²** | ***p*-value** |  | **Element** | **Coefficient β** | **R²** | ***p*-value** |
| Cu | –4.5 | 0.51 | <0.001 |  | Cu | –92 | 0.60 | <0.001 |
| Zn | –7.8 | 0.42 | <0.01 |  | Zn | –99 | 0.61 | <0.001 |
| As | –0.4 | 0.69 | <0.001 |  | As | –4.8 | 0.66 | <0.001 |
| Se | –0.2 | 0.74 | <0.001 |  | Se | –1.6 | 0.63 | <0.001 |
| Cd | –0.03 | 0.34 | <0.05 |  | Cd | –1.8 | 0.72 | <0.001 |
| Pb | –0.01 | 0.07 | >0.05 |  | Pb | –2.2 | 0.49 | <0.001 |
| Ti | 1.3 | 0.57 | <0.001 |  | Ti | 7.9 | 0.10 | >0.05 |
| Co | –0.10 | 0.11 | >0.05 |  | Co | –1.5 | 0.42 | <0.01 |
| Sr | –15 | 0.58 | <0.001 |  | Sr | 0.08 | 0.01 | >0.05 |
| Tl | –0.02 | 0.18 | >0.05 |  | Tl | –0.3 | 0.11 | >0.05 |
| La | 0.05 | 0.47 | <0.01 |  | La | –0.03 | 0.006 | >0.05 |
| Ce | 0.08 | 0.42 | <0.01 |  | Ce | 0.06 | 0.004 | >0.05 |

Table S11. Concentration (mean ± standard deviation, sd; n= 3-10; nmol/g dry weight, dw) of trace elements (TEs) including technology-critical (TCEs) elements measured in walleye’s liver in the six studied lakes. Letters that are different indicate significant differences between means (Kruskal-Wallis, followed by Dunn honestly post hoc test on ranks, *p* < 0.05).

| **Lake**  **(CODE)** | **Osisko**  **(N1)** | | **Rouyn**  **(N2)** | **Dufault**  **(N3)** | **Vaudray**  **(I1)** | **Opasatica**  **(F1)** | **Dufay**  **(F2)** |
| --- | --- | --- | --- | --- | --- | --- | --- |
| **TE** | | |  |  |  |  |  |
| **Cu** | 428 ± 234 ^a^ | | 445 ± 275 ^ab^ | 316 ± 112 ^a^ | 138 ± 133 ^b^ | 129 ± 27 ^b^ | 176 ± 30 ^ab^ |
| **As** | 5 ± 4 ^a^ | | 1 ± 0.30 ^b^ | 3 ± 0.91 ^ab^ | 4.50 ± 2.60 ^ab^ | 3.90 ± 2 ^ab^ | 2.90 ± 0.70 ^ab^ |
| **Zn** | 1967 ± 453 ^a^ | | 1462 ± 290 ^ab^ | 1611 ± 484 ^ab^ | 1246 ± 442 ^b^ | 1242 ± 152 ^b^ | 1869 ± 238 ^a^ |
| **Cd** | 111 ± 44 ^ab^ | | 27 ± 7 ^ac^ | 137 ± 37 ^b^ | 79 ± 41 ^abc^ | 26 ± 11 ^c^ | 87 ± 62 ^abc^ |
| **Fe** | 31525 ± 8331 ^a^ | | 9984 ± 4682 ^ab^ | 9208 ± 4555 ^b^ | 6176 ± 3192 ^b^ | 6003 ± 3179 ^b^ | 8843 ± 2481 ^b^ |
| **Se** | 239 ± 87 ^a^ | | 177 ± 77 ^abc^ | 171 ± 39 ^ab^ | 53 ± 20 ^cd^ | 38 ± 5 ^d^ | 79 ± 15 ^bcd^ |
| **Pb** | 0.27 ± 0.18 ^ab^ | | 0.11 ± 0.03 ^ab^ | 0.46 ± 0.40 ^a^ | 0.17 ± 0.21 ^b^ | 0.15 ± 0.25 ^b^ | 0.30 ± 0.19 ^ab^ |
| **TCE** | | |  |  |  |  |  |
| **Ti** | | 8 ± 1.75 ^a^ | 9 ± 0.36 ^a^ | 7 ± 1.30 ^ab^ | 4 ± 1 ^b^ | 8.8 ± 1.19 ^a^ | 8 ± 2 ^a^ |
| **Co** | 22 ± 10 ^ab^ | | 12 ± 3 ^ab^ | 20 ± 13 ^ab^ | 15 ± 7 ^ab^ | 10 ± 4 ^a^ | 31 ± 16 ^b^ |
| **Sr** | 5 ± 4 ^ab^ | | 2.63 ± 1.54 ^a^ | 9 ± 12 ^a^ | 5 ± 3 ^ab^ | 2 ± 0.87 ^a^ | 22 ± 20 ^b^ |
| **Tl** | 0.60 ± 0.18 ^ab^ | | 5 ± 3 ^a^ | 1 ± 0.37 ^a^ | 0.28 ± 0.14 ^b^ | 0.71 ± 0.19 ^ab^ | 0.53 ± 0.12 ^b^ |
| **La** | 0.04 ± 0.008 ^a^ | | 0.08 ± 0.04 ^abc^ | 0.08 ± 0.02 ^ab^ | 0.12 ± 0.03 ^bc^ | 0.37 ± 0.46 ^bc^ | 0.86 ± 0.29 ^c^ |
| **Ce** | 0.03 ± 0.01^a^ | | 0.08 ± 0.02 ^abc^ | 0.06 ± 0.01^ab^ | 0.07 ± 0.02 ^ab^ | 0.11 ± 0.04 ^bc^ | 0.63 ± 0.24 ^c^ |

Table S12. Concentration (mean ± standard deviation; n= 3-10; nmol/g dry weight, dw) of trace elements (TEs) including technology-critical (TCEs) elements measured in yellow perch’s liver in the six studied lakes. Letters that are different indicate significant differences between means (Kruskal-Wallis, followed by Dunn honestly post hoc test on ranks, *p* < 0.05).

| **Lake**  **(CODE)** | **Osisko**  **(N1)** | | **Rouyn**  **(N2)** | **Dufault**  **(N3)** | **Vaudray**  **(I1)** | **Opasatica**  **(F1)** | **Dufay**  **(F2)** |
| --- | --- | --- | --- | --- | --- | --- | --- |
| **TE** | | |  |  |  |  |  |
| **Cu** | 1781 ± 1682 ^a^ | | 550 ± 336 ^ab^ | 319 ± 139 ^ab^ | 645 ± 469 ^ab^ | 121 ± 3 ^b^ | 788 ± 590 ^ab^ |
| **As** | 3 ± 1 ^ab^ | | 3 ± 0.83 ^abc^ | 1 ± 0.45 ^c^ | 2.70 ± 0.92 ^abc^ | 8 ± 4 ^a^ | 2 ± 1 ^bc^ |
| **Zn** | 2914 ± 702 ^a^ | | 1617 ± 191 ^abc^ | 1501 ± 164 ^b^ | 2238 ± 427 ^abc^ | 1434 ± 16 ^bc^ | 2631 ± 842 ^ac^ |
| **Cd** | 200 ± 163 ^ab^ | | 61 ± 21 ^ab^ | 153 ± 44 ^a^ | 199 ± 137 ^a^ | 43 ± 3 ^ab^ | 43 ± 69 ^b^ |
| **Fe** | 46965 ± 33847 ^a^ | | 16340 ± 3278 ^ab^ | 8261 ± 3179 ^b^ | 8687 ± 4742 ^b^ | 10078 ± 20 ^ab^ | 91479 ± 81196 ^a^ |
| **Se** | 231 ± 51 ^ab^ | | 303 ± 49 ^a^ | 256 ± 40 ^a^ | 99 ± 33 ^c^ | 71 ± 1 ^bc^ | 94 ± 36 ^c^ |
| **Pb** | 0.78 ± 1.21 ^ab^ | | 0.39 ± 0.13 ^abc^ | 0.98 ± 0.35 ^ac^ | 2 ± 0.66 ^c^ | 0.12 ± 0.004 ^b^ | 0.50 ± 0.29 ^ab^ |
| **TCE** | | |  |  |  |  |  |
| **Ti** | | 9 ± 2 ^abc^ | 26 ± 10 ^a^ | 7 ± 1 ^bc^ | 9 ± 5 ^abc^ | 5 ± 0.36 ^b^ | 11 ± 3 ^ac^ |
| **Co** | 11 ± 6 ^a^ | | 11 ± 2 ^ab^ | 8 ± 4 ^a^ | 9 ± 4 ^a^ | 6.8 ± 0.47 ^a^ | 101 ± 61 ^b^ |
| **Sr** | 27 ± 22 ^a^ | | 1 ± 0.25 ^ab^ | 0.8 ± 0.12 ^b^ | 12 ± 15 ^a^ | 1 ± 0.22 ^ab^ | 9 ± 6 ^a^ |
| **Tl** | 0.18 ± 0.08 ^ab^ | | 22 ± 5 ^c^ | 1 ± 0.38 ^c^ | 0.58 ± 0.32 ^abc^ | 1 ± 0.11 ^ac^ | 0.18 ± 0.15 ^b^ |
| **La** | 0.12 ± 0.09 ^ab^ | | 0.07 ± 0.03 ^ab^ | 0.04 ± 0.03 ^a^ | 0.20 ± 0.12 ^b^ | 0.22 ± 0.02 ^ab^ | 0.31 ± 0.22 ^b^ |
| **Ce** | 0.16 ± 0.15 ^ab^ | | 0.08 ± 0.04 ^ab^ | 0.04 ± 0.03 ^a^ | 0.19 ± 0.15 ^b^ | 0.17 ± 0.04 ^ab^ | 0.38 ± 0.40 ^b^ |

Table S13. Summary of linear regression analyses of trace element concentrations in walleye and yellow perch liver as a function of distance from the Horne smelter. Regression coefficients (β), R² values, and significance levels are reported. Data from all sampled lakes were included for each element (n=3-10/lake).

| **Walleye** |  |  |  |  | **Yellow perch** |  |  |  |
| --- | --- | --- | --- | --- | --- | --- | --- | --- |
| **Element** | **Coefficient β** | **R²** | ***p*-value** |  | **Element** | **Coefficient β** | **R²** | ***p-*value** |
| Cu | -7 | 0.362 | <0.001 |  | Cu | -13 | 0.04 | >0.05 |
| Zn | -5.4 | 0.033 | >0.05 |  | Zn | 4.5 | 0.009 | >0.05 |
| As | -0.01 | 0.007 | >0.05 |  | As | 0.003 | 0.001 | >0.05 |
| Se | -4.3 | 0.600 | <0.001 |  | Se | -4.6 | 0.7 | <0.001 |
| Cd | -1.1 | 0.104 | <0.05 |  | Cd | -2.2 | 0.08 | >0.05 |
| Pb | -0.003 | 0.022 | >0.05 |  | Pb | 0.006 | 0.01 | >0.05 |
| Ti | -0.02 | 0.016 | >0.05 |  | Ti | -0.01 | 0.001 | >0.05 |
| Co | 0.09 | 0.012 | >0.05 |  | Co | 1.6 | 0.3 | <0.001 |
| Sr | 0.2 | 0.089 | <0.05 |  | Sr | -0.2 | 0.02 | >0.05 |
| Tl | -0.03 | 0.123 | <0.05 |  | Tl | -0.09 | 0.06 | >0.05 |
| La | 0.01 | 0.463 | <0.001 |  | La | 0.006 | 0.3 | <0.001 |
| Ce | 0.01 | 0.424 | <0.001 |  | Ce | 0.006 | 0.2 | <0.01 |

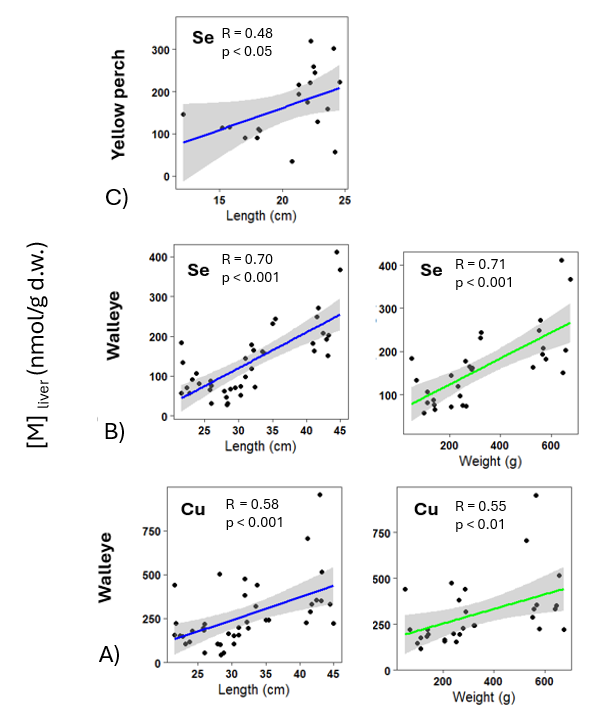

Figure S1. Relationship between trace element concentrations in fish liver (nmol/g dry weight, dw) and biometric variables (length and weight, in centimeters and grams, respectively) across six studied lakes.
**(A)** Cu concentrations in walleye liver vs length (left) and weight (right);
**(B)** Se concentrations in walleye liver vs length (left) and weight (right);
**(C)** Se concentrations in yellow perch liver vs length.

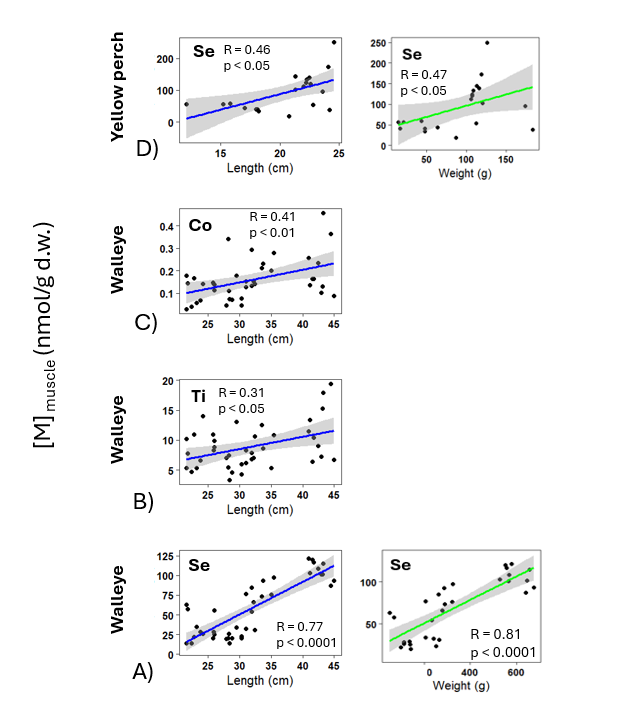

Figure S2. Relationship between trace element concentrations in fish muscle (nmol/g dry weight, dw) and biometric variables (length and weight, in centimeters and grams, respectively) across six studied lakes.
**(A)** Se concentrations in walleye muscle vs length (left) and weight (right);

**(B)** Ti concentrations in walleye muscle vs length (left);

**(C)** Co concentrations in walleye muscle vs length (left);

**(D)** Se concentrations in yellow perch muscle vs length (left) and weight (right).

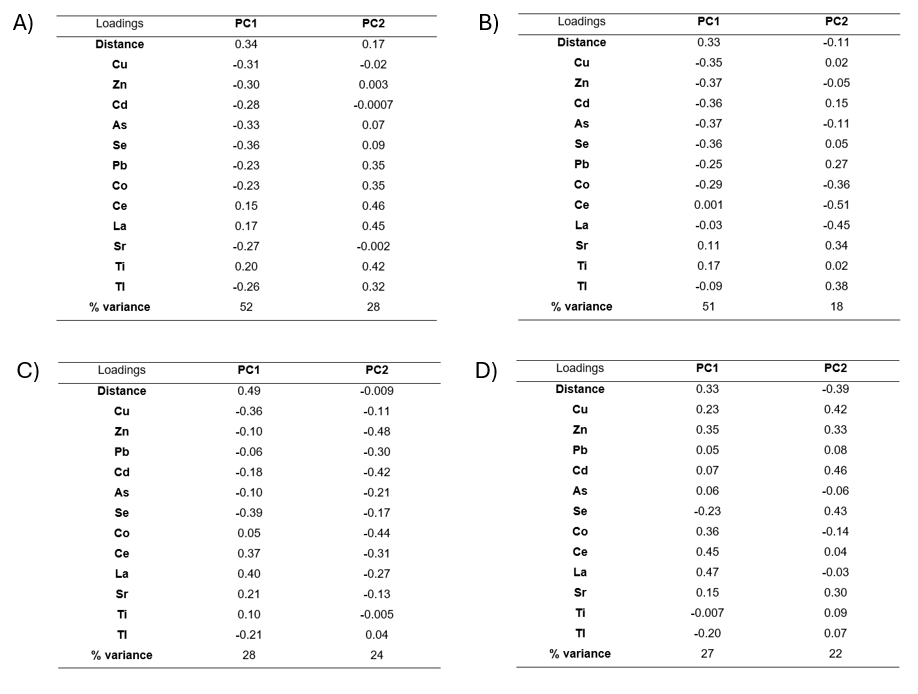

Figure S3. PCA loadings of the different variables in A) water, B) sediment, C) walleye liver, and D) yellow perch liver corresponding to PC1 and PC2. These results complement the PCA analyses presented in Figures 6 and 7 of the main article.

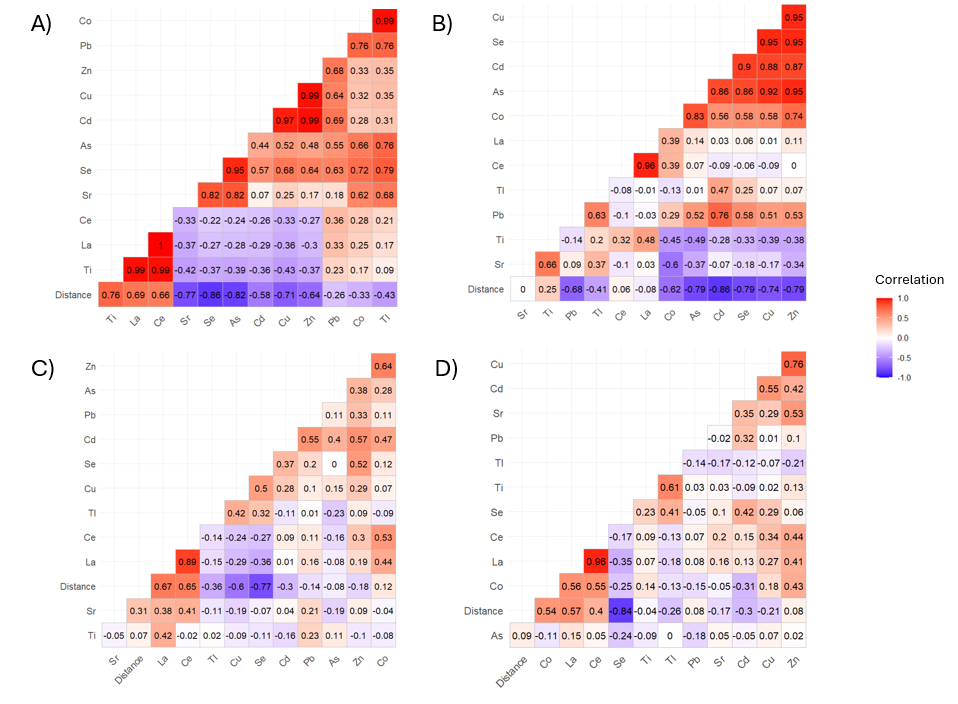

Figure S4. Spearman correlation coefficients for relationships between trace element concentrations in A) water, B) sediment, C) walleye liver, and D) yellow perch liver samples and the distance from the Horne smelter. These results complement the PCA analyses presented in Figures 6 and 7 of the main article. All correlations are significant, *p* < 0.05, n =18-49.

CCME. (2007). *Canadian environmental quality guidelines.* Canadian Council of Ministers of the Environment. https://www.ccme.ca/en

Sneller, F. E. C., Kalf, D. F., Weltje, J., & Van Wezel, A. P. (2000). *Maximum Permissible Concentrations and Negligible Concentrations for Rare Earth Elements (REEs)*. National Istitute of Public Health and the Environment.

Van Son, M. (1994). *De bepaling van het gehalte aan “exotische metalen” in sediment- en water monster.* [TNO--report TNO-MW-R94/319]. National Istitute of Public Health and the Environment.

Wetzel, R. G., & Likens, G. E. (2000). *Limnological Analyses*. Springer New York. https://doi.org/10.1007/978-1-4757-3250-4
